# Localized disruption of the presynaptic endoplasmic reticulum in *Atlastin* mutants

**DOI:** 10.1101/2023.09.01.555994

**Authors:** Mónica C. Quiñones-Frías, Dina M. Ocken, Avital A. Rodal

## Abstract

The endoplasmic reticulum (ER) extends throughout neurons and regulates many neuronal functions, including neurite outgrowth, neurotransmission, and synaptic plasticity. Mutations in proteins that control ER shape are linked to the neurodegenerative disorder Hereditary Spastic Paraplegia (HSP), yet the ultrastructure and dynamics of neuronal ER remain largely unexplored, especially at presynaptic terminals. Using super-resolution and live imaging in *Drosophila melanogaster* larval motor neurons, we investigated ER structure at presynaptic terminals of wild-type animals and null mutants of the HSP-linked gene, *Atlastin*, which encodes an ER-shaping protein. Previous studies reported diffuse localization of an ER luminal marker at *Atlastin* null mutant presynaptic terminals, which was attributed to ER fragmentation. However, using an ER membrane marker, we found that *Atlastin* mutant ER forms robust ER networks with only mild defects in structure and dynamics, indicating that the primary defect is functional rather than architectural. We demonstrate that *Atlastin* mutants progressively displace a luminal ER protein reporter to the cytosol during larval development, specifically at synapses, while this reporter remains correctly localized in cell bodies, axons, and muscles. This synapse-specific displacement phenotype, previously unreported in non-neuronal cells, emphasizes the importance of studying neurons to understand HSP pathogenesis.

## Introduction

The endoplasmic reticulum (ER) is a continuous organelle that extends to the periphery of neurons and regulates many neuronal functions, including neurite outgrowth, neurotransmission, and synaptic plasticity (1–6). Structurally, this essential organelle consists of two domains: ER sheets (enriched in cell bodies) and ER tubules (abundant in dendrites and axons) (7,8). Recent studies have revealed even finer levels of organization, with increasing evidence suggesting structural and functional specialization of neuronal ER tubules. For instance, different regions of the neuron show distinct tubule features, including varying densities of ER-organelle contacts (8), specialized ER-endosome contacts that regulate lysosome size in cell bodies (9), and unique ER ladder structures in developing axons and dendrites (10,11). These regional specializations underscore the importance of examining ER structure and function across all neuronal regions. Studying the cell biology of neuronal ER is particularly important because mutations in ER-shaping genes cause neurodegenerative disorders such as Hereditary Spastic Paraplegia (HSP) and Charcot Marie Tooth Disease, indicating that ER structure is uniquely essential for neuronal physiology (7,12,13). However, the cell biology and ultrastructure of the neuronal ER, particularly at presynaptic terminals, have been under-investigated.

One essential ER tubule-shaping protein is Atlastin, a GTPase that regulates the homotypic fusion and tethering of ER tubules to form 3-way junctions (14). Mutations in *Atlastin 1,* one of three human *Atlastin* genes, underlie one of the most common forms of autosomal dominant HSP, highlighting the importance of examining its neuron-specific functions in regulating ER structure (15). In various experimental systems, *Atlastin* mutants exhibit numerous neuronal phenotypes, including defects in synaptic vesicle cycling and trafficking of neuronal cargoes (16), synapse development (16–18), and axon regeneration (19). However, significant gaps remain in our understanding of how Atlastin-dependent changes in the structure and dynamics of neuronal ER lead to these functional defects. For example, using electron microscopy, null mutants of the single *Drosophila Atlastin* homolog were found to exhibit shorter ER tubules in their motor neuron cell bodies (20) and a diffuse localization of an ER lumen marker in their presynaptic terminals, attributed to extensive ER fragmentation (21). In *C. elegans, Atlastin* null mutant ER networks in neuronal dendrites were prone to retraction, resulting in disrupted microtubule structures and mitochondrial fission (2). Finally, a ladder-like ER expansion was observed in spinal axons in a mouse model of Atlastin-dependent HSP (22). In contrast to these neuronal findings, knockdown of Atlastin in Hela or Cos-7 cells leads to long and unbranched ER tubules (23–25). Thus, loss of Atlastin causes distinct ER phenotypes that depend on several factors, including cell type, the region of the neuron examined, and the reporter used to visualize the ER. To elucidate the functions of Atlastin, comprehensive imaging studies of ER organization across different regions of the neuron are needed. This is best done in neurons *in vivo*, in their native context and making functional synaptic connections, since HSP is a progressive, adult-onset disease of long motor axons whose relevant ER defects may depend on synaptic function.

Another critical gap is our limited understanding of neuronal ER dynamics. High-resolution imaging in non-neuronal cells has revealed transitions from ER tubules to ER sheets (26,27), ER fragmentation (28), tubule extension and retraction (29–31), and alterations in the number of 3-way junctions (32). These dynamics have been linked to calcium signaling (27,33) and ER functions such as endosomal and mitochondrial fission (29–31). In neurons, ER rearrangements have been suggested to alter calcium compartmentalization (34), local protein synthesis (35,36), organelle contacts (37), and transport within the ER (38). However, the types and functions of neuronal ER dynamics, particularly in presynaptic terminals, remain largely unknown due to the challenges of imaging ER networks over time in small compartments. New insights into the mechanisms that regulate ER structure and ER dynamics in neurons are crucial for understanding why and how mutations in ER shaping proteins, such as Atlastin, predominantly affect the nervous system.

Conventional light microscopy, commonly used in studies of neuronal ER structure, lacks the resolution necessary to visualize individual ER tubules in small structures, such as presynaptic terminals. The ER is highly sensitive to fixation, and live imaging experiments in neurons *in vivo* have been conducted on upright microscopes using water dipping objectives with a typical axial resolution limit of >300 nm, which cannot distinguish the densely packed ER tubules at presynaptic terminals (3,8,21,28,39–42). Electron microscopy offers higher resolution, but cannot be used in live samples and has typically been limited to thin 2D sampling (in which it is difficult to distinguish ER cross-sections from synaptic vesicles) (8,20,22). Here we addressed these technical limitations by examining the structure and dynamics of presynaptic ER at the *Drosophila* neuromuscular junction (NMJ) of wild-type and *Atlastin* mutants using *in vivo* super-resolution microscopy.

## Results

High resolution imaging of the ER in *Drosophila* motor neuron cell bodies, axons, and presynaptic terminals.

We created a reusable microscopy imaging slide to capture live super-resolution images or movies of neuronal ER in *Drosophila* larval fillets through glass coverslips, utilizing high numerical aperture oil objectives on a super-resolution Airyscan microscope (**Figure 1A**). Our system provides an axial resolution of ∼170 nm after image processing (43). We imaged larval fillets using this imaging slide on both inverted and upright microscopes within ∼45 min of dissection, a time frame in which this preparation maintains muscle potential and the capacity for evoked neurotransmitter release, even once the axon is severed from the cell body (44).

**Figure 1.**
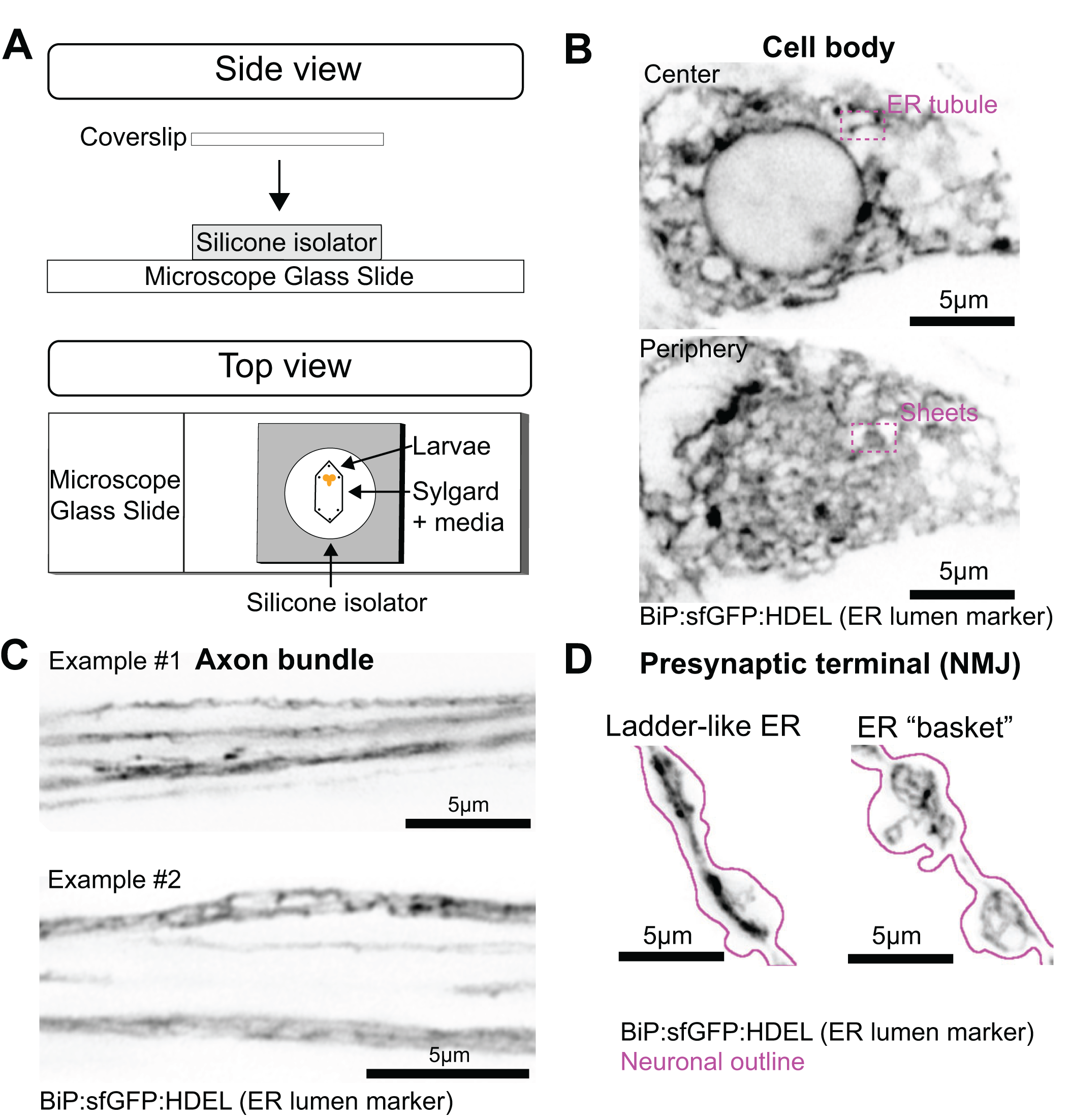
*In vivo* imaging of ER networks in *Drosophila* motor neurons. (**A**) Schematic of live imaging chamber, consisting of a glass microscope slide with a mounted silicone isolator filled with Sylgard. Small metal pins were used to stretch the larvae. Finally, a coverslip was placed on top of the larval fillet for imaging with an oil objective on upright or inverted microscopes. (**B**) A representative single slice of an Airyscan microscopy Z-stack of the center (top) or periphery (bottom) of the cell body of a motor neuron expressing the luminal ER marker BiP:sfGFP:HDEL. Pink dashed boxes indicate ER tubules and sheets. (**C**) Maximum projections of an Airyscan microscope Z-stack of two different motor neuron axon bundles expressing the luminal ER marker BiP:sfGFP:HDEL. (**D**) Representative maximum projection of an Airyscan microscopy Z-stack of a motor neuron presynaptic terminal expressing the luminal ER marker BiP:sfGFP:HDEL. Magenta outline highlights the neuronal boundary.

To study the live dynamics of the ER, we used Vglut-GAL4 to express UAS-driven markers of the ER lumen and ER membrane in *Drosophila* 3^rd^ instar larval motor neurons. The ER lumen was visualized by expressing sfGFP flanked by an N-terminal signal sequence of the ER chaperone BiP (MKLCILLAVVAFVGLSLGRS) and a C-terminal HDEL retention signal (hereafter referred to as BiP:sfGFP:HDEL) (21). To visualize ER membranes, we expressed an UAS line containing tdTomato-tagged full-length Sec61β (hereafter referred to as tdTomato: Sec61β). Our improved imaging approach enabled more precise visualization of ER structures such as tubules and sheets in cell bodies, and tubules in axon bundles and presynaptic terminals (**Figures 1B–D**). In axon bundles, ER tubules were long, and we sometimes observed a ladder-like structure consisting of pairs of interconnected ER tubules joined by rungs or cross-bridges, similar to structures previously described in developing axons of cultured mammalian neurons and *Drosophila* dendrites (10,11) (**Figure 1C**). At presynaptic terminals, ER tubules in some boutons appeared in a highly interconnected basket-like network, while other boutons were traversed by several parallel ER tubules, similar to the ladder-like structures observed in axons (**Figure 1D**). Thus, using this improved imaging approach, we achieved high-resolution visualization of individual ER tubules within networks of *Drosophila* presynaptic terminals, allowing for a detailed analysis of their structure and dynamics over time.

We next characterized ER dynamics in *en passant* and terminal boutons (**Figure 2A**). Compared to *en passant* boutons, terminal boutons exhibit distinct cytoskeletal organization and signaling, which could regulate the dynamics of the ER differently (45–47). We collected Z-stacks of single boutons expressing BiP:sfGFP:HDEL for 90 sec at 0.73 sec/stack (**Figure 2B**, **Movie S1–S3**) or tdTomato:Sec61β for 40 sec at 0.92 sec/stack (**Figure 2C, Movie S4–S6**). Both markers were distributed unevenly across the ER network (**Figure 2B–2C**). In particular, the luminal ER marker exhibited numerous discrete round foci that moved within ER tubules with fast dynamics (<1 µm*/*sec) (**Figure 2B**). These foci were less frequently observed with the ER membrane marker (**Figure 2C**, **bottom**), consistent with previous findings that proteins in the ER lumen and ER membrane differentially cluster and move within the ER (48,49). Conversely, static regions of high intensity (which we saw for both markers) may reflect stacked ER tubules (50).

**Figure 2.**
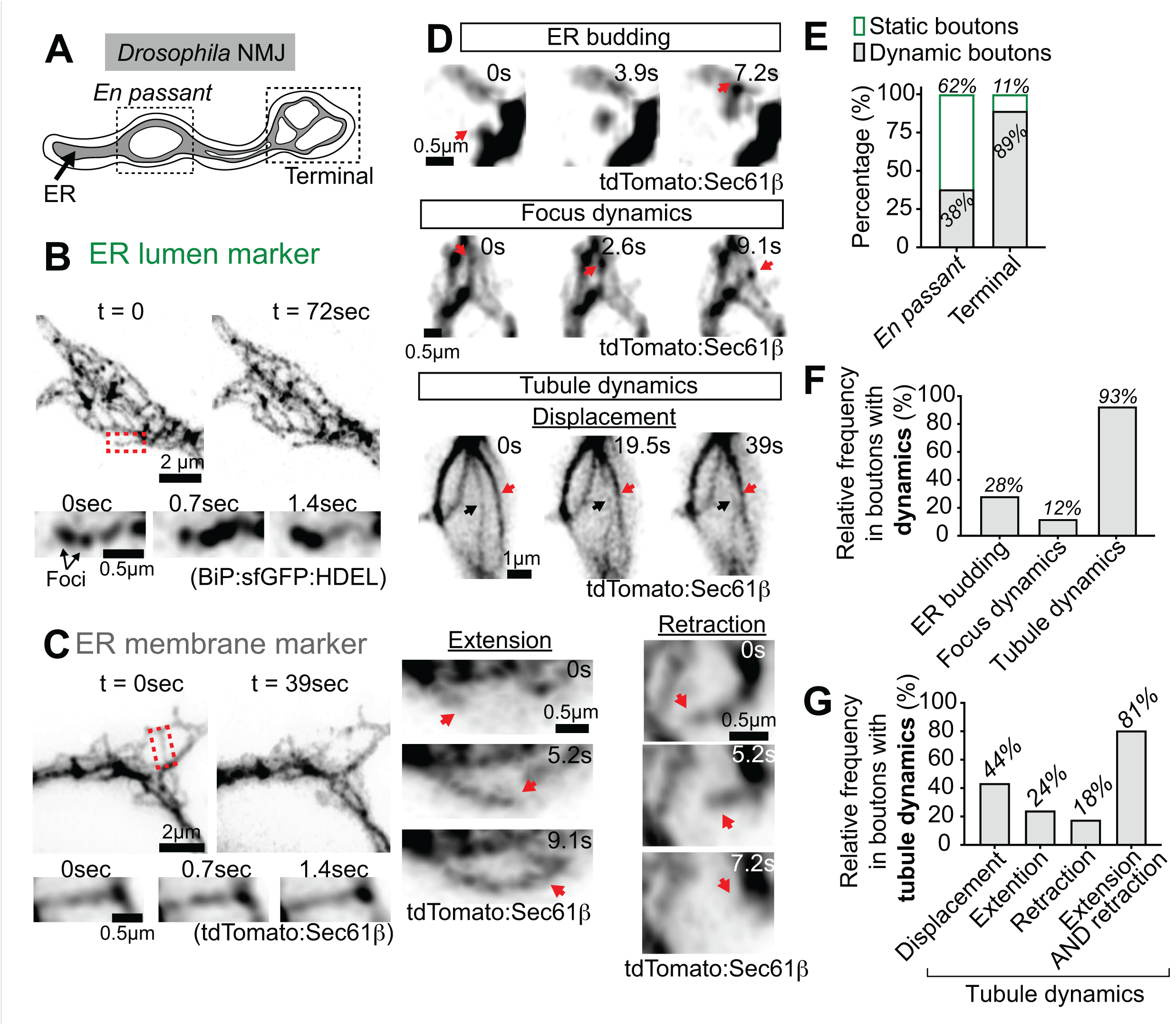
Dynamics of luminal and membrane ER markers at presynaptic terminals. (**A**) Diagram of a presynaptic terminal of *Drosophila* motor neurons highlighting *en passant* and terminal boutons with the ER in grey. (**B, top**) Maximum intensity projection of Airyscan Z-stacks from the beginning (t=0) and the end (t=72 sec) of a time-lapse movie (**Movie S1**) of a single wild-type bouton expressing the luminal ER marker BiP:sfGFP:HDEL. Other examples can be found in **Movies S2–S3**. (**B, bottom**) Magnification of the region in the dotted red box of dynamics observed from 0 sec to 1.4 sec of the movie. (**C**, **top**) Maximum intensity projection of Airyscan Z-stacks from the beginning (t=0) and the end (t=39 sec) of a time-lapse movie (**Movie S4**) of a single wild-type bouton expressing the ER membrane marker tdTomato:Sec61β. Other examples can be found in **Movies S5–S6**. (**C, bottom**) Magnification of the region in the dotted red box of dynamics observed from 0 sec to 1.4 sec of the movie. (**D**) Presynaptic terminals exhibiting ER budding (**Movie S7**), focus dynamics (**Movie S8**), and various types of ER tubule dynamics (ER tubule displacement (**Movie S9**), tubule extension (**Movie S10**), and tubule retraction (**Movie S11**)). Red arrows highlight types of dynamic events. For tubule displacement, black arrows indicate a stationary ER tubule that remained in the same position throughout the time series. Note that the red and black arrows converge over time as the mobile tubule moves toward the stationary one. (**E**) Qualitative categorization of the percentage of *en passant* and terminal boutons that are dynamic or static, expressing the ER membrane marker tdTomato:Sec61β. (**F**) Qualitative categorization of the percentage of dynamic terminal boutons expressing the ER membrane marker tdTomato:Sec61β that exhibit ER budding, foci dynamics, or ER tubule dynamics. (**G**) Qualitative categorization of the percentage of dynamic terminal boutons expressing the ER membrane marker tdTomato:Sec61β that exhibit tubule displacement, tubule extension, tubule retraction and both tubule extension and retraction. The same control dataset used in E-G was used in Figure 5. Detailed genotype, replicate, and statistical information can be found in Supplementary Table 1.

We focused on the ER membrane marker to define dynamic events, since its labeling pattern had fewer confounding foci. Analysis revealed three distinct types of ER dynamics: (1) ER budding events, in which a fragment of ER dissociates from the network; (2) Focus dynamics, in which round foci move within the ER network; and (3) ER tubule dynamics, including overall displacement of the tubule in the XY plane, as well as extension and retraction of the tubule (**Figure 2D, Movies S7–11**). We imaged boutons for 40 sec at 0.92 sec intervals to capture ER dynamics over this observation period. Boutons were qualitatively categorized as “static” if we observed no detectable changes in ER network structure throughout the entire 40 sec imaging session, or “dynamic” if we observed at least one of the three defined dynamic events during this time window. On many occasions, we observed multiple types of ER dynamics within the same bouton. We found that terminal boutons are highly dynamic (89%) while *en passant* boutons were moderately dynamic (38%) (**Figure 2E**). This difference in dynamics suggests that the ER is regulated differently in different regions of presynaptic terminals and could have specialized functions.

Next, we characterized the frequency of the three main types of ER network dynamics at terminal boutons, where we observed the most remodeling. Among terminal boutons, 93% exhibited tubule movements, making this the most common type of ER remodeling (**Figure 2F**). Note that individual NMJs could exhibit multiple ER dynamics simultaneously, resulting in overlapping categories that sum to more than 100%. Less frequent events included ER budding (28% of boutons) and moving foci (12% of boutons). We further classified tubule movements into four categories: displacement, extension, retraction, and extension followed by retraction. The most common category was extension followed by retraction, observed in 81% of terminal boutons (**Figure 2G**). Simple displacement occurred in 44% of boutons, 24% showed extension alone, and 18% showed retraction alone. These results demonstrate our ability to capture diverse ER behaviors at high resolution in live *Drosophila* motor neurons.

### Presynaptic terminals of *Atlastin* mutants form robust ER networks

To investigate the relationship between ER structure and function at synapses, we examined mutants of Atlastin, a GTPase that regulates ER tubule fusion. The diverse neuronal phenotypes in *Atlastin* mutants, including defects in synaptic growth, vesicle cycling, cargo trafficking, and axon regeneration, make it an excellent model for understanding how ER architecture impacts cellular function (16–19,51). We first assessed the architecture of ER networks in *Atlastin* mutants by visualizing the ER membrane marker tdTomato:Sec61β in transheterozygotes of an *Atlastin* null allele (*atl*^2^) and a deficiency removing the *Atlastin* locus to avoid second-site effects (17). These *Atlastin* null mutants (hereafter referred to as *Atlastin* mutants) were mainly pharate lethal and showed increased satellite boutons at larval NMJs (**Figure 3A–B**), indicating defects in synaptic development as previously reported (16,17). Surprisingly, we found that tdTomato:Sec61β labeled intact ER networks in *Atlastin* mutant synapses (**Figure 3A**, **Figure 3—figure supplement 1**), unlike the previously reported diffuse localization of a luminal marker in this mutant (21). We quantified the tdTomato:Sec61β signal and found an increase in the coefficient of variation (CoV) and lower fluorescence levels compared to controls (**Figure 3C–D**). Lower fluorescence could indicate thinner tubules or reduced protein content in ER tubules. A higher CoV indicates an uneven distribution of tdTomato:Sec61β within the presynaptic terminal, with some areas showing higher concentrations than others (in contrast to the uniform, diffuse signal expected from fragmentation). These data indicate that ER networks remain largely intact in *Atlastin* mutant presynaptic terminals, and that the previously observed functional defects likely result from subtle changes in ER structure rather than the complete fragmentation of the ER network. Lower levels of ER membrane proteins have also been observed in other ER-shaping mutants, suggesting this is a common feature (41). To test this, we examined mutants of *Reticulon1*, another HSP-linked ER shaping factor, and found similarly reduced tdTomato:Sec61β levels compared to controls (**Figure 3E–F**). This finding suggests that ER shaping mutants share phenotypes, likely because they contribute to some of the same functions.

**Figure 3.**
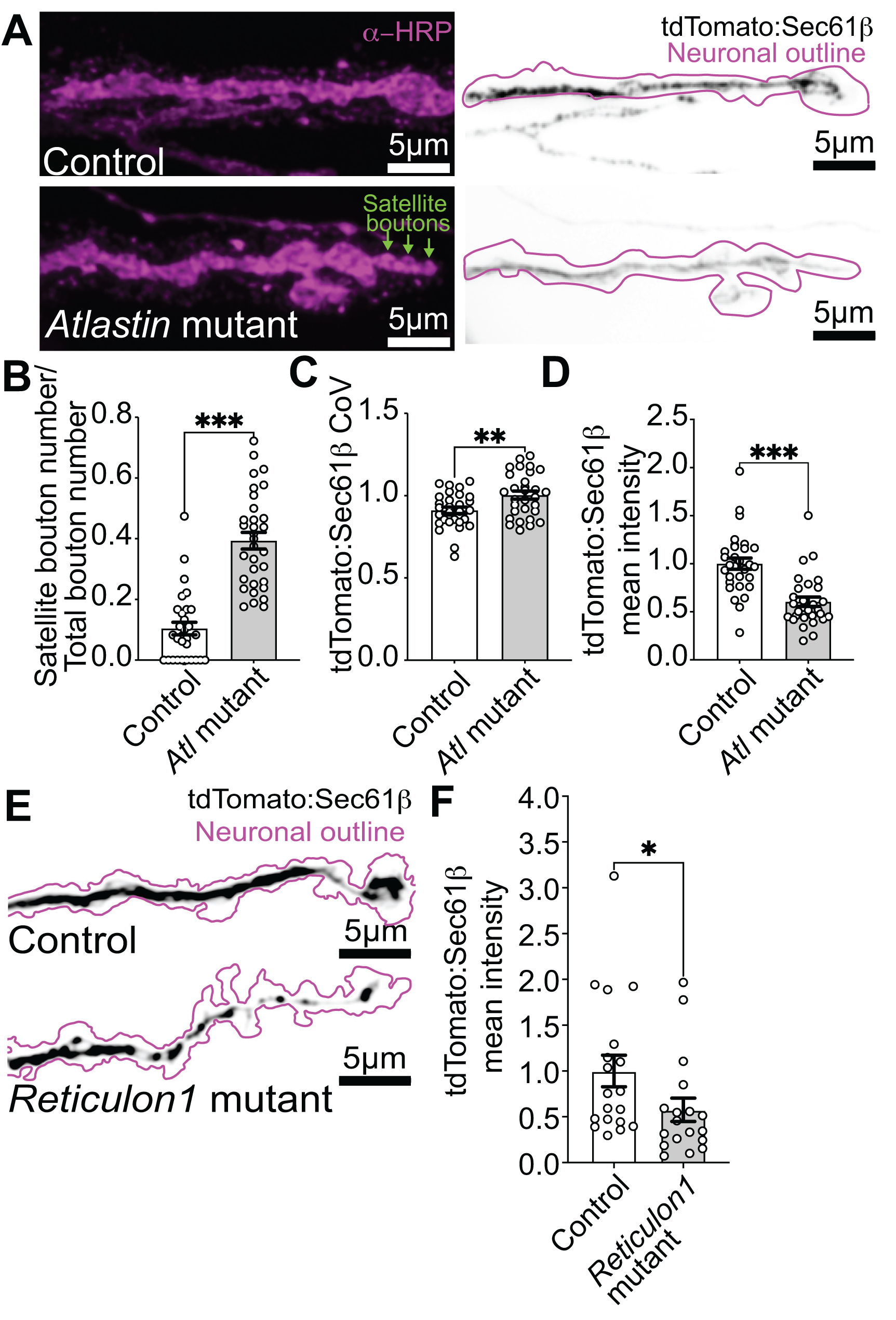
*Atlastin* and *Reticulon1* mutants have lower levels of the ER membrane tdTomato:Sec61β at presynaptic terminals. (**A**) Representative maximum intensity projection of a spinning disk confocal Z-stack from fixed larvae expressing the ER membrane marker tdTomato:Sec61β (left) and neuronal membranes labeled by α−HRP (right) in control (**top**) and *Atlastin* mutants (**bottom**). Green arrows highlight a string of satellite boutons. (**B**) Quantification of satellite boutons normalized to total bouton number in control and *Atlastin* mutants. (**C**) Coefficient of variation (CoV) of the ER membrane marker tdTomato:Sec61ββ . (**D**) Quantification of intensity of the ER membrane marker tdTomato:Sec61ββ, normalized to wild-type control. (**E**) Representative maximum intensity projection of a SoRa Z-stack from live larvae expressing the ER membrane marker tdTomato:Sec61ββ (black) in control (**top**) and *Reticulon1* mutants (**bottom**). (**F**) Quantification of the intensity of the ER membrane marker tdTomato:Sec61ββ , normalized to wild-type control. (**B-D, F**). Detailed genotype, replicate, and statistical information can be found in Supplementary Table 1.

**Figure 3—figure supplement 1.**
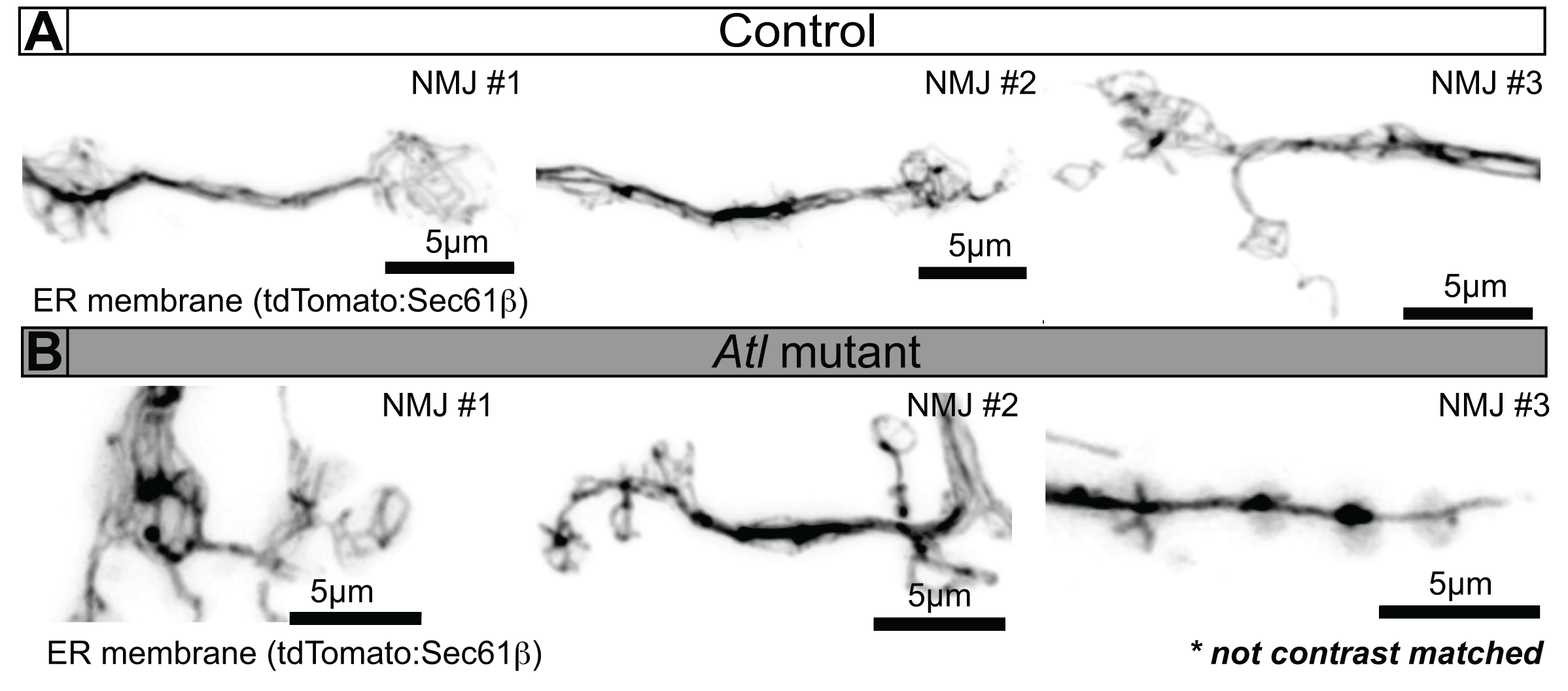
*Atlastin* mutants expressing the ER membrane tdTomato:Sec61ββ **exhibit robust ER networks at presynaptic terminals.** (**A, B**) Representative maximum projection of a Z-stack acquired live by Airyscan microscopy of control and *Atlastin* mutant NMJs expressing the ER membrane marker tdTomato:Sec61ββ at presynaptic terminals. These images were not contrast-matched to facilitate visualization of ER structures across all genotypes (See Figure 3D).

We measured the levels of tdTomato:Sec61β in various motor neuron regions to determine if its reduced levels are specific to presynaptic terminals or a cell-wide phenotype (**Figure 4**). Using Vglut-GAL4 to label motor neurons, we measured tdTomato:Sec61β in clusters of cell bodies in the ventral ganglion and axon bundles extending towards body-wall muscles. *Atlastin* mutants had significantly increased tdTomato:Sec61β intensity levels in axon bundles and unchanged levels in clusters of cell bodies (**Figure 4C–D**). We also found that the volume occupied by tdTomato:Sec61β-labeled structures was significantly increased in axon bundles and decreased in clusters of cell body in *Atlastin* mutants (**Figure 4E–F**). Finally, we observed a significant decrease in the CoV of the tdTomato:Sec61β signal in both axon bundles and cell body clusters, suggesting less concentrated ER structures (the opposite of what we observed at presynaptic terminals) (**Figure 4G–H**). Overall, we observed distinct changes in cell body clusters, axon bundles, and presynaptic terminals, indicating that the ER has neuronal region-specific functions regulated by Atlastin. These changes in tdTomato:Sec61β1 distribution could reflect alterations in ER structure or ER membrane protein density and are notably distinct from the previously reported diffuse localization of an ER luminal marker (21).

**Figure 4.**
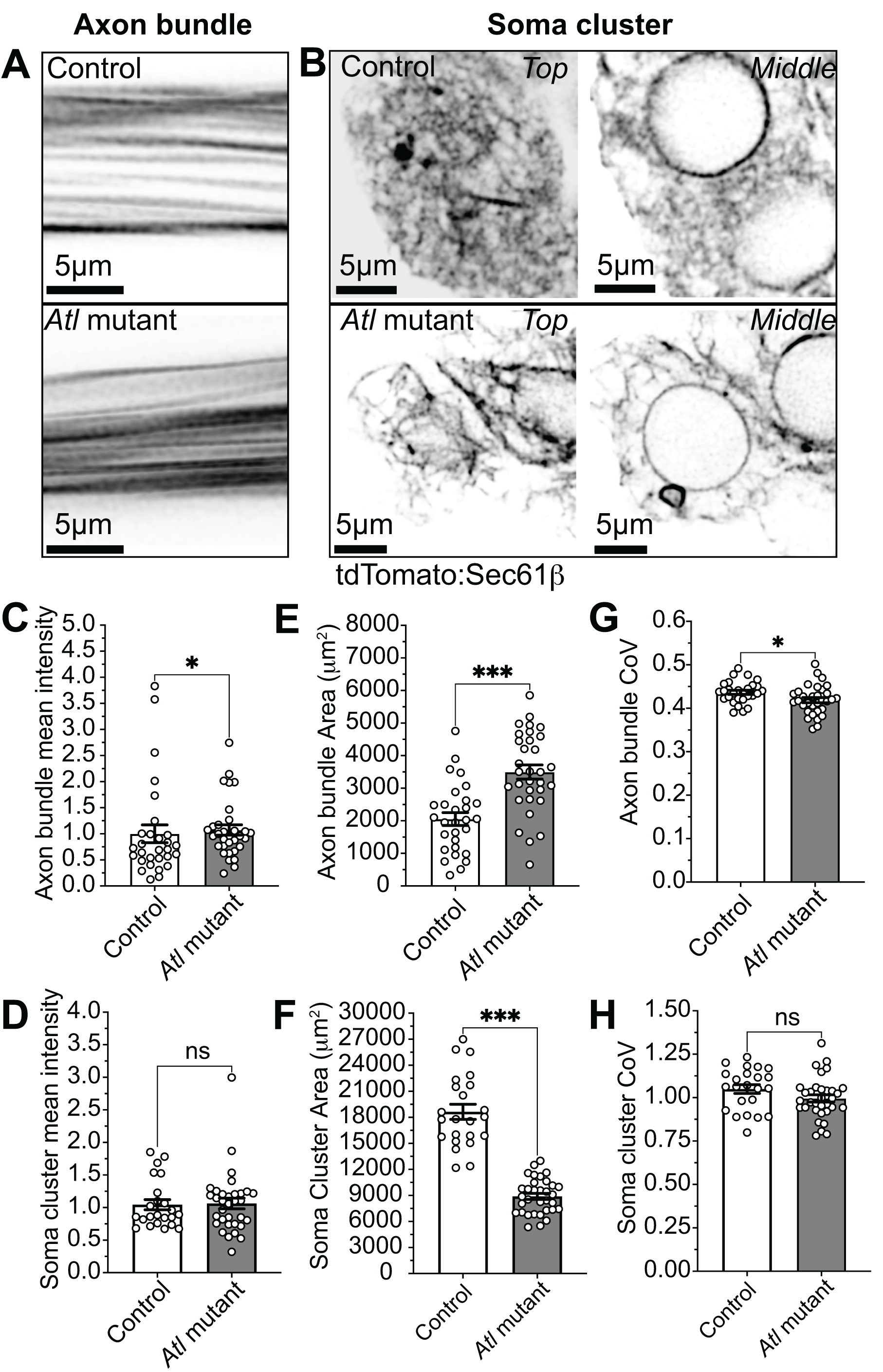
Atlastin regulates ER membrane marker distribution in a compartment-specific manner. (**A**) Representative middle slice of axon bundles in control (top) and *Atlastin* mutants (bottom) expressing tdTomato:Sec61β. (**B**) Representative top and middle slices of somas in control (top) and *Atlastin* mutants (bottom) expressing tdTomato:Sec61β. (**C,D**) Quantification of the intensity of the ER membrane marker tdTomato:Sec61β. (**E,F**) Quantification of the area of the ER membrane marker tdTomato:Sec61β. (**G,H**) Coefficient of variation (CoV) of the ER membrane marker tdTomato:Sec61β. Detailed genotype, replicate, and statistical information can be found in Supplementary Table 1.

### *Atlastin* mutants show mild defects in synaptic ER dynamics

We examined the role of Atlastin in regulating ER network dynamics, given its function as an ER-shaping protein. We captured time-lapse images of the ER labeled with tdTomato:Sec61β in *en passant* and terminal boutons at 0.92 sec/stack (**Figure 5A, Movies S12–S19**). We blinded movies from controls and *Atlastin* mutants to qualitatively categorize boutons as “dynamic” or “static”, as described above (**Figure 2E**). Compared to the 89% of terminal boutons in control animals that exhibited dynamics (same dataset used in **Figure 2B, E**), we found a small but significant reduction in dynamic boutons in *Atlastin* mutants (76%), with a concomitant increase in the fraction of non-dynamic boutons (24%) (**Figure 5B**). In *en passant* boutons, we observed similar ER dynamics in controls (62% static/38% dynamic) and *Atlastin* mutants (69% static/31% dynamic). These findings indicate that *Atlastin* mutants exhibit only mild defects in ER dynamics, specifically at terminal boutons, with no significant changes in *en passant* boutons. This suggests that Atlastin is not critical for the dynamics of ER networks at presynaptic terminals.

**Figure 5.**
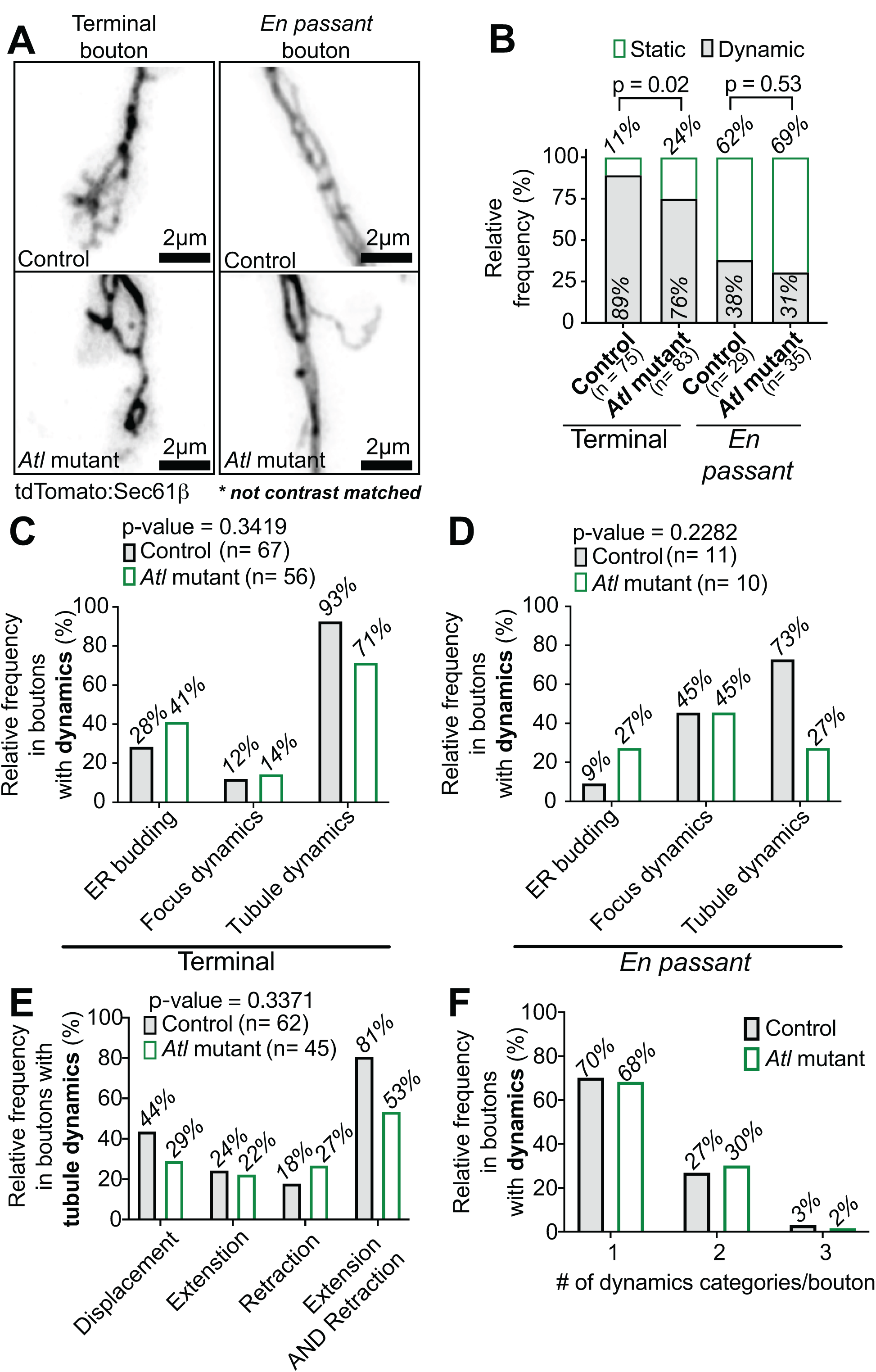
*Atlastin* mutants have mild ER dynamics phenotypes. (**A**) Representative images acquired with an Airyscan microscope of terminal (left) and *en passant* (right) boutons in controls (top) and *Atlastin* mutants (bottom) (**Movies S12–19**). Control and *Atlastin* mutant images are not contrast-matched, enabling visualization of ER structures across all genotypes. (**B**) Percentage of dynamic versus static boutons in controls and *Atlastin* mutants expressing the ER membrane marker tdTomato:Sec61β, analyzed separately for terminal and *en passant* boutons. (**C**) Distribution of ER dynamic types (budding, foci dynamics, or ER tubule dynamics) in dynamic terminal boutons expressing tdTomato:Sec61β. (**D**) Distribution of ER dynamic types in dynamic *en passant* boutons expressing tdTomato:Sec61β. (**E**) Qualitative categorization of the percentage of dynamic terminal boutons in controls and *Atlastin* mutants expressing the ER membrane marker tdTomato:Sec61β that exhibit tubule displacement, tubule extension, tubule retraction, and both tubule extension and retraction. (**F**) Qualitative categorization of the number of dynamic categories per terminal bouton in control and *Atlastin* mutants. Data from control animals were also used in Figure 2. Detailed genotype, replicate, and statistical information can be found in Supplementary Table 1.

We next performed an analysis of the types of ER dynamics (displacement, extension, retraction, and extension followed by retraction) in *Atlastin* mutants. Surprisingly, despite Atlastin being the sole homolog in *Drosophila* and a key player in ER membrane fusion, we found only modest changes in ER dynamics. While we observed slight shifts in the relative frequencies of different dynamic events, none of these changes reached statistical significance (**Figure 5C–E**). We also counted the number of dynamic events per bouton to assess its dynamic capacity, and *Atlastin* mutants remained unaffected (**Figure 5F**). The subtlety of these phenotypes suggests either substantial functional redundancy with other ER-shaping proteins or compensatory mechanisms that maintain ER organization and dynamics in neurons upon loss of *Atlastin*. Moreover, our findings suggest that the numerous functional defects previously reported at *Atlastin* mutant synapses may arise from defects in ER function rather than gross changes in ER structure or dynamics.

### *Atlastin* mutant presynaptic terminals exhibit distinct BiP:sfGFP:HDEL and tdTomato:Sec61β distributions

We found that *Atlastin* mutants have robust ER networks, as indicated by the ER membrane marker tdTomato:Sec61β. However, these results stand in sharp contrast to a prior study that examined ER networks using the luminal marker BiP:sfGFP:HDEL, in which a diffuse distribution was interpreted as ER fragmentation (21). To further explore this discrepancy, we used Airyscan microscopy to re-examine BiP:sfGFP:HDEL in *Atlastin* mutants (**Figure 6A–B**). Unlike the strict localization of this lumenal marker to the ER network in control synapses, we reproduced the observation that a significant fraction of BiP:sfGFP:HDEL was diffusely localized in *Atlastin* mutant synapses, even at the resolution of Airyscan microscopy. Furthermore, the marker filled the interiors of boutons and extended to their peripheries, suggesting that it may be mislocalized to the cytosol rather than within ER fragments.

**Figure 6.**
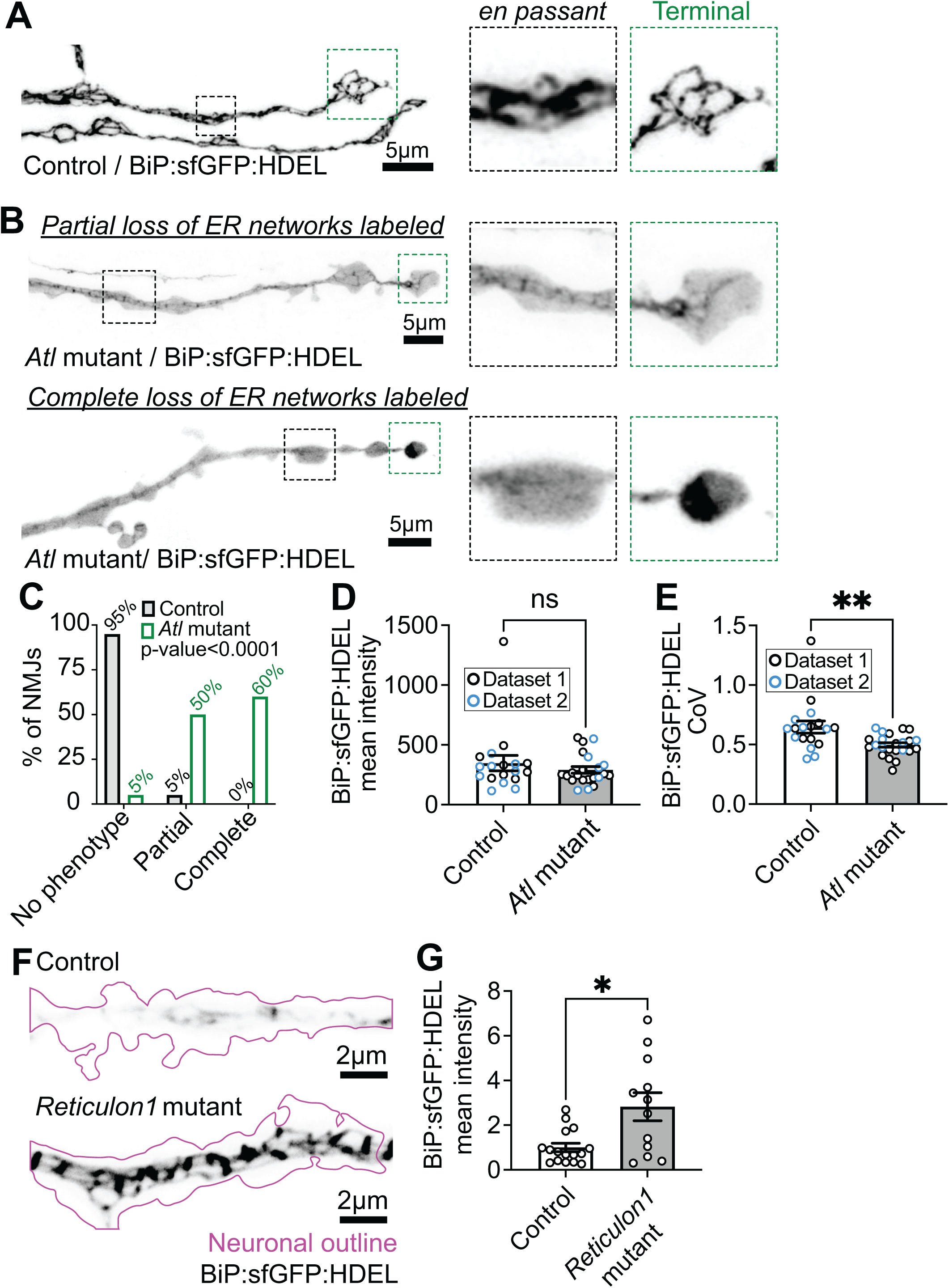
Luminal ER proteins are redistributed to a diffuse pattern in *Atlastin* mutants. Maximum projection of an Airyscan Z-stack of control (**A**) and *Atlastin* mutant (**B**) NMJs expressing the ER lumen marker BiP:sfGFP:HDEL. (**B**) *Atlastin* mutants exhibit two distinct phenotypes at presynaptic terminals that were categorized as partial loss (top) or complete loss (bottom). Control and *Atlastin* mutant images are contrast matched. (**C**) Qualitative categorization of presynaptic terminals in control and *Atlastin* mutants expressing BiP:sfGFP:HDEL that exhibit “no phenotype”, “partial”, or “complete” loss of the luminal marker localized to the ER. (**D**) Quantification of the intensity of the luminal ER marker BiP:sfGFP:HDEL. (**E**) Coefficient of variation (CoV) of the luminal ER marker BiP:sfGFP:HDEL (**F**) Maximum projection a SoRa Z-stack of control (**top**) and *Reticulon1* mutant (**bottom**) NMJs expressing the ER lumen marker BiP:sfGFP:HDEL. (**G**) Quantification of intensity of the luminal ER marker BiP:sfGFP:HDEL. Detailed genotype, replicate, and statistical information can be found in Supplementary Table 1.

**Figure 6—figure supplement 1.**
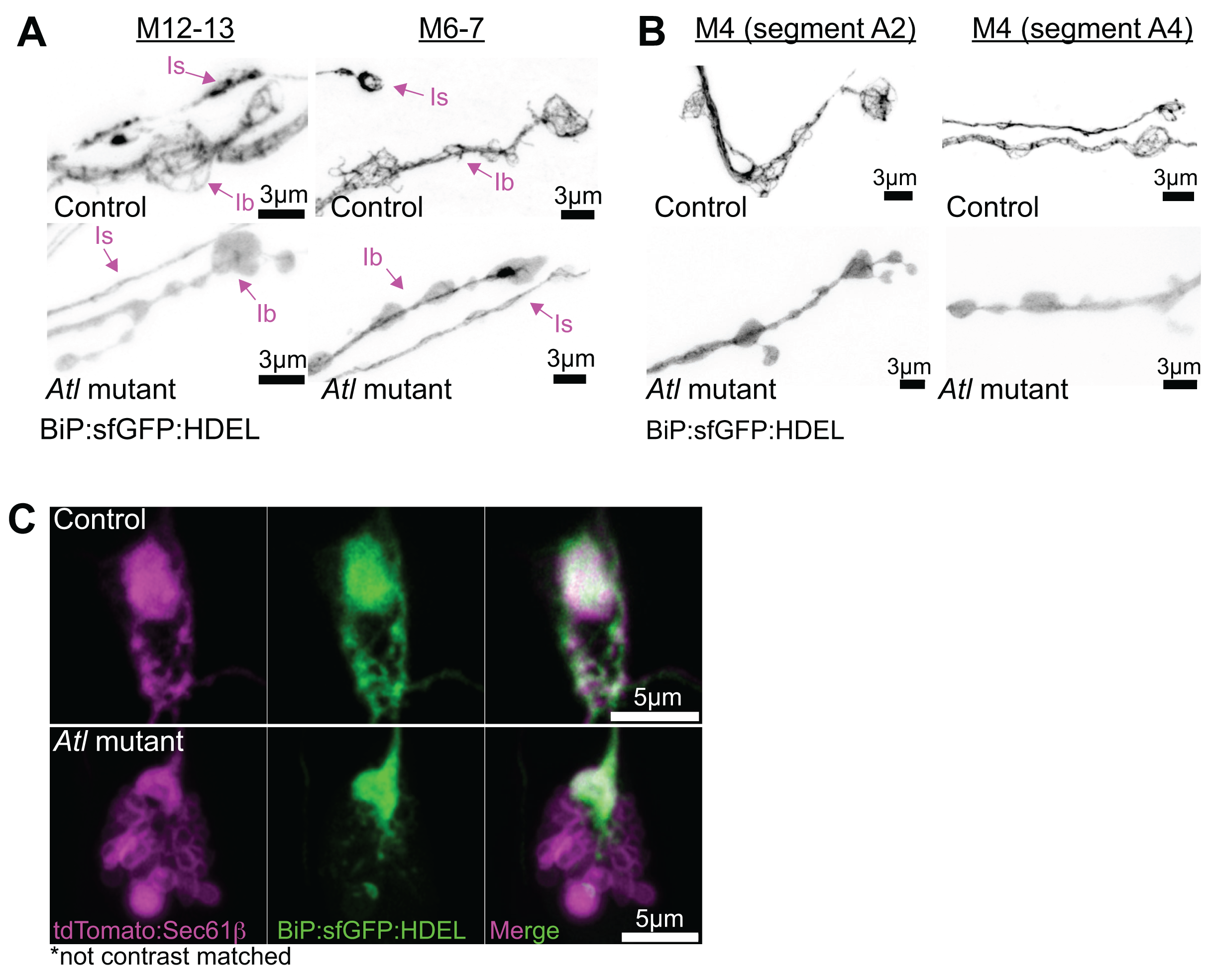
Examples of Is and Ib motor neurons in various muscles and abdominal segments of 3^rd^ instar *Atlastin* mutant larvae exhibiting cytosolic displacement of BiP:sfGFP:HDEL. Maximum projection of a live Airyscan Z-stack from larvae expressing the ER lumen marker BiP:sfGFP:HDEL at muscles 12-13 (**A, left**), muscles 6-7 (**A, right**), muscle 4 in segment A2 (**B, left**) and segment A4 (**B, right**). Co-expression of tdTomato:Sec61β and BiP:sfGFP:HDEL severely disrupts ER at NMJs in *Atlastin* mutants. Representative single slice images acquired with an Airyscan microscope of (**C**) control (top) and *Atlastin* mutant (bottom) NMJs showing tdTomato:Sec61β (left), BiP:sfGFP:HDEL (middle), and merge (right).

We identified two distinct ER network phenotypes in *Atlastin* mutants expressing BiP:sfGFP:HDEL: “Partial loss” NMJs retained both diffuse signal and identifiable ER network structures, while “Complete loss” NMJs showed no visible ER network structures. Note that the “Complete loss” phenotype in *Atlastin* mutants reflects the absence of detectable luminal marker signal in organized ER structures, not the complete absence of ER membranes, as demonstrated by our ER membrane marker tdTomato:Sec61β results. Blinded categorization revealed that 50% of *Atlastin* mutant NMJs exhibited partial loss, 60% complete loss, and 5% no phenotype (**Figure 6C**). In contrast, controls showed 5% partial loss, 0% complete loss and 95% no phenotype. We note that the sum of these percentages exceeds 100% because one NMJ exhibited multiple phenotypes: one branch had a complete loss, while the other branch had no phenotype. These phenotypes were counted separately.

*Atlastin* mutant presynaptic terminals showed similar mean intensities of the luminal ER marker compared to controls, suggesting that redistribution does not affect overall marker levels (**Figure 6D**). However, we observed a significant reduction in BiP:sfGFP:HDEL CoV in *Atlastin* mutants due to the redistribution of the luminal marker (**Figure 6E**). This phenotype was consistent across all motor neurons examined, including both anterior (segment A2) and posterior (segment A5) muscle segments, in Is (phasic) and Ib (tonic) motor neuron subtypes, and across multiple body wall muscles (**Figure 6—figure supplement 1**). Interestingly, this phenotype is not universal among ER-shaping mutants; we found that BiP:sfGFP:HDEL expressed in *Reticulon1* mutants localized robustly to ER networks, and that its levels were higher than in controls (**Figure 6F–G**). Finally, we were unable to assess co-localization between BiP:sfGFP:HDEL and tdTomato:Sec61β in *Atlastin* mutants due to poor animal survival and grossly defective NMJ morphologies, characterized by large vesicles (**Figure 6—figure supplement 1**). However, combined with our earlier finding that the ER membrane marker tdTomato:Sec61β remains intact when expressed alone, these results suggest that the luminal ER marker is mislocalized away from the ER, and that this phenotype is specific to *Atlastin* mutants.

### The displacement of luminal ER proteins in *Atlastin* mutants is specific to presynaptic terminals

Next, we asked whether the luminal ER marker displacement phenotype is cell-autonomous to neurons by knocking down neuronal Atlastin using GAL4-driven UAS-*Atl* RNAi, compared to mCherry RNAi controls. We observed a phenotype similar to that of the null *Atl* mutant at presynaptic terminals, indicating that luminal displacement results from loss of neuronal Atlastin (**Figure 7A**). Further quantification showed that neuronal Atlastin knockdown, compared to the mCherry RNAi control, showed no change in BiP:sfGFP:HDEL mean intensity but a significant decrease in CoV (**Figure 7B–C**), similar to *Atl* null mutants. This finding suggests Atlastin function is required in neurons to maintain proteins in the ER lumen at presynaptic terminals.

**Figure 7.**
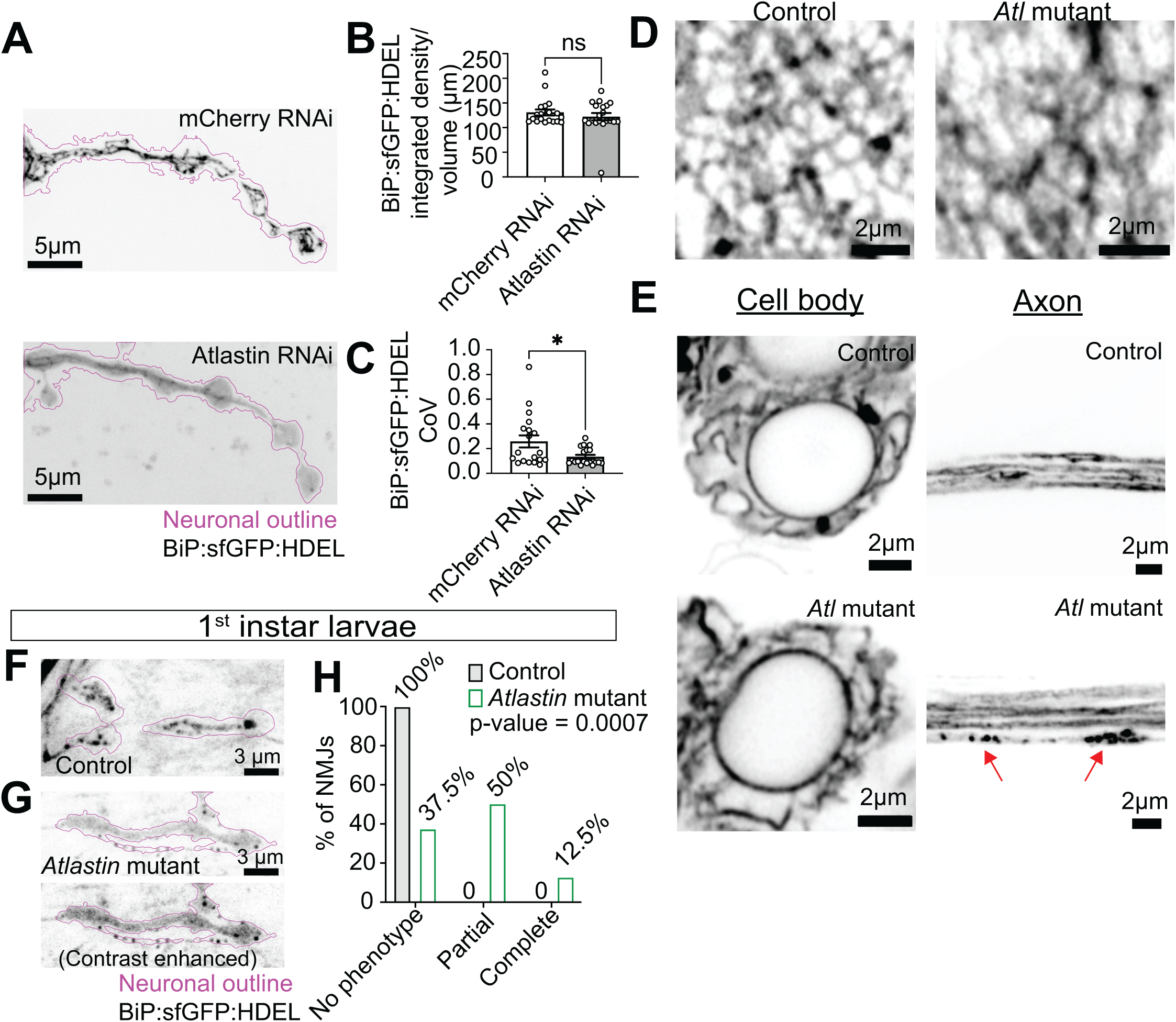
Luminal ER proteins are progressively displaced only at presynaptic terminals of *Atlastin* mutants. (**A**) Representative maximum projection live SoRa Z-stack of BiP:sfGFP:HDEL of control and neuronal Atlastin RNAi. (**B**) Quantification of the intensity of the luminal ER marker BiP:sfGFP:HDEL and (**C**) CoV of the luminal ER marker BiP:sfGFP:HDEL in neurons of control and neuronal *Atlastin* RNAi. (**D**) A single slice of body wall muscles acquired with live Airyscan microscopy from larvae of control and *Atlastin* mutants expressing the ER lumen marker BiP:sfGFP:HDEL with BG57-GAL4 (body wall muscles). (**E**) Maximum projection of a Z-stack acquired with live Airyscan microscopy of a single slice of motor neuron cell bodies (left) and motor neuron axon bundles (right). (**F, G**) Maximum intensity projections of a live Airyscan Z-stack of 1^st^ instar larvae expressing the ER lumen marker BiP:sfGFP:HDEL (black; neuronal outline in magenta) in control (F) and *Atlastin* mutants (G). (**H**) Qualitative categorization of presynaptic terminals in control and *Atlastin* mutant 1^st^ instar larvae expressing BiP:sfGFP:HDEL that exhibit “no phenotype”, “partial”, or “complete” loss of the luminal marker localized to the ER. Detailed genotype, replicate, and statistical information can be found in Supplementary Table 1.

We next examined other cell types and neuronal regions to determine whether the luminal ER marker displacement phenotype is specific to neurons and presynaptic terminals. We examined the ER in body wall muscles of *Atlastin* mutants by overexpressing the BiP:sfGFP:HDEL with the muscle driver BG57-Gal4. We did not find the luminal ER marker displaced to the cytosol (**Figure 7D**). However, as noted in earlier studies (17), we observed a decrease in network complexity. Next, we examined the cell body and axons of *Atlastin* null mutants to determine if the displacement of the luminal ER marker was a global or local phenotype within neurons. As previously reported (21), we did not observe a diffuse signal for the luminal ER marker in cell bodies or distal axons (within ∼30µm of the synapse) (**Figure 7E**). However, we did observe ER fragmentation in some axons. These results indicate that the displacement of BiP:sfGFP:HDEL in *Atlastin* mutants is specific to presynaptic terminals.

### The luminal ER phenotype in *Atlastin* mutants is progressive during larval development

We reasoned that the BiP:sfGFP:HDEL displacement phenotype might progress developmentally, similar to progressive neuronal dysfunction in Atlastin-linked HSP patients and progressive motor deficits in *Drosophila Atlastin* mutants (16,52–54). To test this hypothesis, we examined 1^st^ instar larvae. We categorized the distribution of BiP:sfGFP:HDEL in control and *Atlastin* mutant live preparations as described for BiP:sfGFP:HDEL (**Figure 6**). Note that in 1^st^ instar larvae, both normal networks in controls and residual networks in *Atlastin* mutants appeared more fragmented than in 3^rd^ instar preparations, likely due to the technical challenges of dissecting these smaller, more delicate specimens. Since ER fragmentation occurred similarly in both genotypes, we could still reliably assess the redistribution of BiP:sfGFP:HDEL as our primary phenotypic readout. We expected that a progressive phenotype would lead to fewer NMJs exhibiting BiP:sfGFP:HDEL displacement and more presynaptic terminals showing no phenotype. Indeed, in 1^st^ instar larvae NMJs, 12.5% showed complete loss, 50% showed partial loss, and 37.5% exhibited no phenotype (**Figure 7F–H**). By contrast, in NMJs of 3^rd^ instar larvae, 60% showed complete loss, 50% showed partial loss, and 5% showed no phenotype (**Figure 6C**). Our results suggest that displacement of the luminal ER marker progresses over time and that this phenotype is initiated early in neuronal development.

### Luminal ER proteins are displaced to the cytosol at presynaptic terminals of *Atlastin* mutants

We hypothesized that the diffuse distribution of BiP:sfGFP:HDEL reflects its mislocalization to the cytosol in presynaptic terminals of *Atlastin* mutants. We co-expressed BiP:sfGFP:HDEL with morphotrap_Int_ (mCherry:CD8:vhhGFP4), a plasma membrane-tethered GFP nanobody fused to mCherry (55). If BiP:sfGFP:HDEL is mislocalized to the cytosol in *Atlastin* mutants, morphotrap_Int_ should sequester it at the plasma membrane through its cytosol-facing GFP-binding nanobody, reducing the diffuse cytoplasmic signal. Indeed, we found that *Atlastin* mutants co-expressing morphotrap_Int_ and BiP:sfGFP:HDEL had a reduced fraction of cytosolic displacement compared to BiP:sfGFP:HDEL alone (**Figure 6C**), with 47% of NMJs with partial loss, 3% with complete loss, and 50% with no phenotype (**Figure 8A–B**). The altered distribution of BiP:sfGFP:HDEL in *Atlastin* mutants is specifically due to the GFP nanobody in morphotrap_Int_, as co-expression with RFP:mCD8, which lacks the nanobody, results in phenotypes similar to BiP:sfGFP:HDEL expression alone (42% partial loss, 50% complete loss, 8% no phenotype; **Figure 8B** vs **Figure 6C**). The shift in distribution induced by the expression of morphotrap_Int_ indicates that it is interacting with BiP:sfGFP:HDEL on the cytosolic face of membranes in *Atlastin* mutants. We note that rather than accumulating at the plasma membrane as expected, the overall levels of BiP:sfGFP:HDEL were reduced, unmasking residual ER networks in *Atlastin* mutants (**Figure 8C**). However, given that this effect depends on the morphotrap_Int_ tool, it is unlikely to reflect a normal degradative process for cytosolic BiP:sfGFP:HDEL. Finally, in wild-type controls, neither morphotrap_Int_ nor RFP:mCD8 affected BiP:sfGFP:HDEL distribution or levels (**Figure 8C**). Together, these results suggest that a substantial fraction of BiP:sfGFP:HDEL in *Atlastin* mutants is mislocalized to the cytosol.

**Figure 8.**
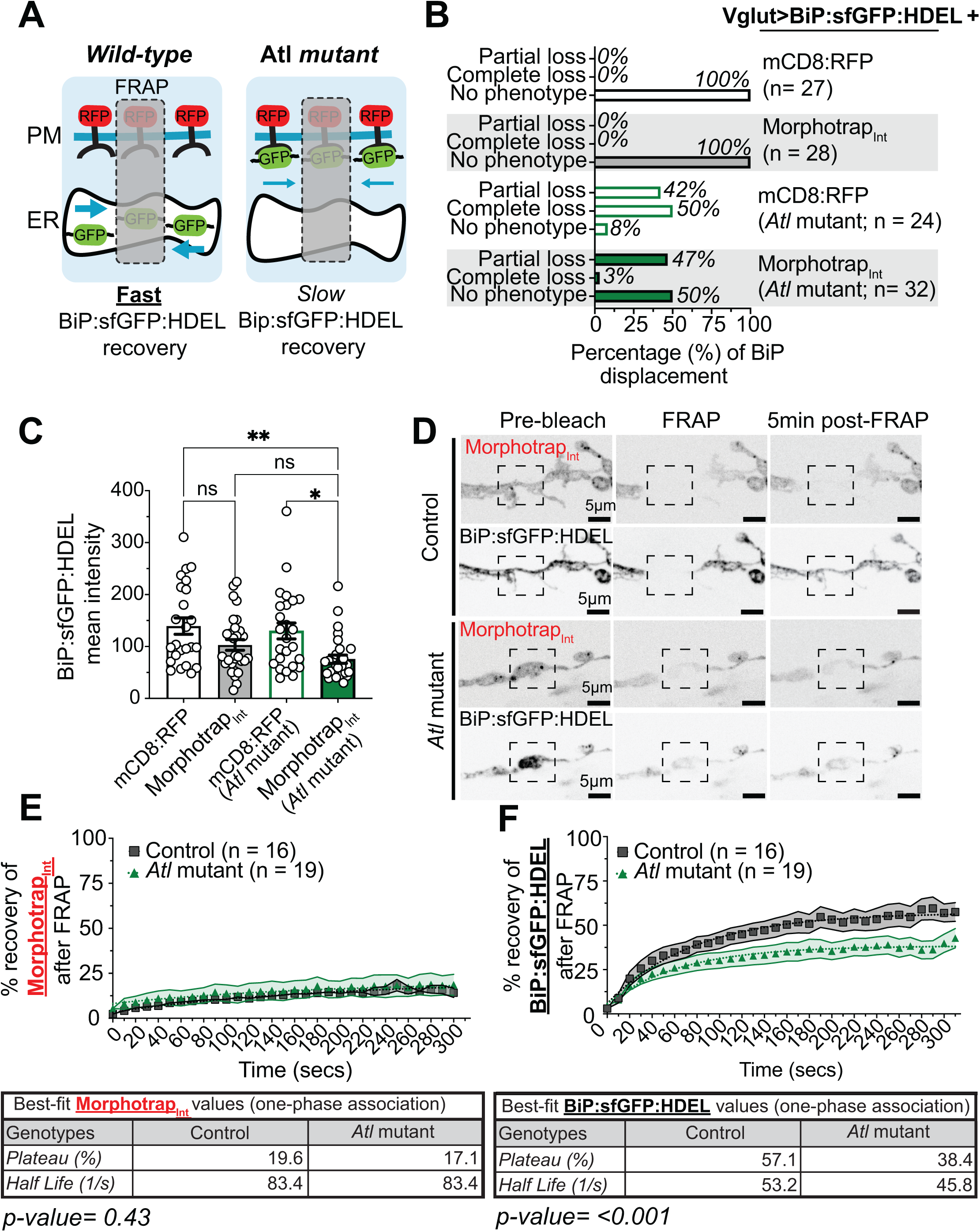
BiP:sfGFP:HDEL is exposed to the cytosol in *Atlastin* mutants. (**A**) Diagram of the recovery of BiP:sfGFP:HDEL in controls and *Atlastin* mutants co-expressing morphotrap_Int_. Blue arrow thickness indicates the rate of recovery. (**B**) Distribution of BiP:sfGFP:HDEL phenotypes (partial loss, complete loss, or no phenotype) in control and *Atlastin* mutant presynaptic terminals co-expressing either mCD8:RFP or morphotrap_Int_. (**C**) Intensity of BiP:sfGFP:HDEL in control and *Atlastin* mutant terminals co-expressing either mCD8:RFP or morphotrap_Int_. (**D**) Representative maximum intensity projections of SoRA Z-stacks from controls (also see **Movies S20–S21**) and *Atlastin* mutants (also see **Movies S22–S23**) co-expressing morphotrap_Int_ and BiP:sfGFP:HDEL pre-bleach, immediately after FRAP and 5 min post-FRAP. The dashed boxes in (D) indicate areas that were photobleached and analyzed for recovery quantification in (E–F). (**E**) Mean recovery of mCD8:RFP in control (black squares) and *Atlastin* mutants (green triangles). (**F**) Mean recovery of BiP:sfGFP:HDEL in control (black squares) and *Atlastin* mutants (green triangles). Table shows non-linear fit one-phase association parameters. s.e.m. are represented by solid lines and shading: controls have black lines with grey shading, *Atlastin* mutants have green lines with light green shading. Detailed genotype, replicate, and statistical information can be found in Supplementary Table 1.

**Figure 8—figure supplement 1.**
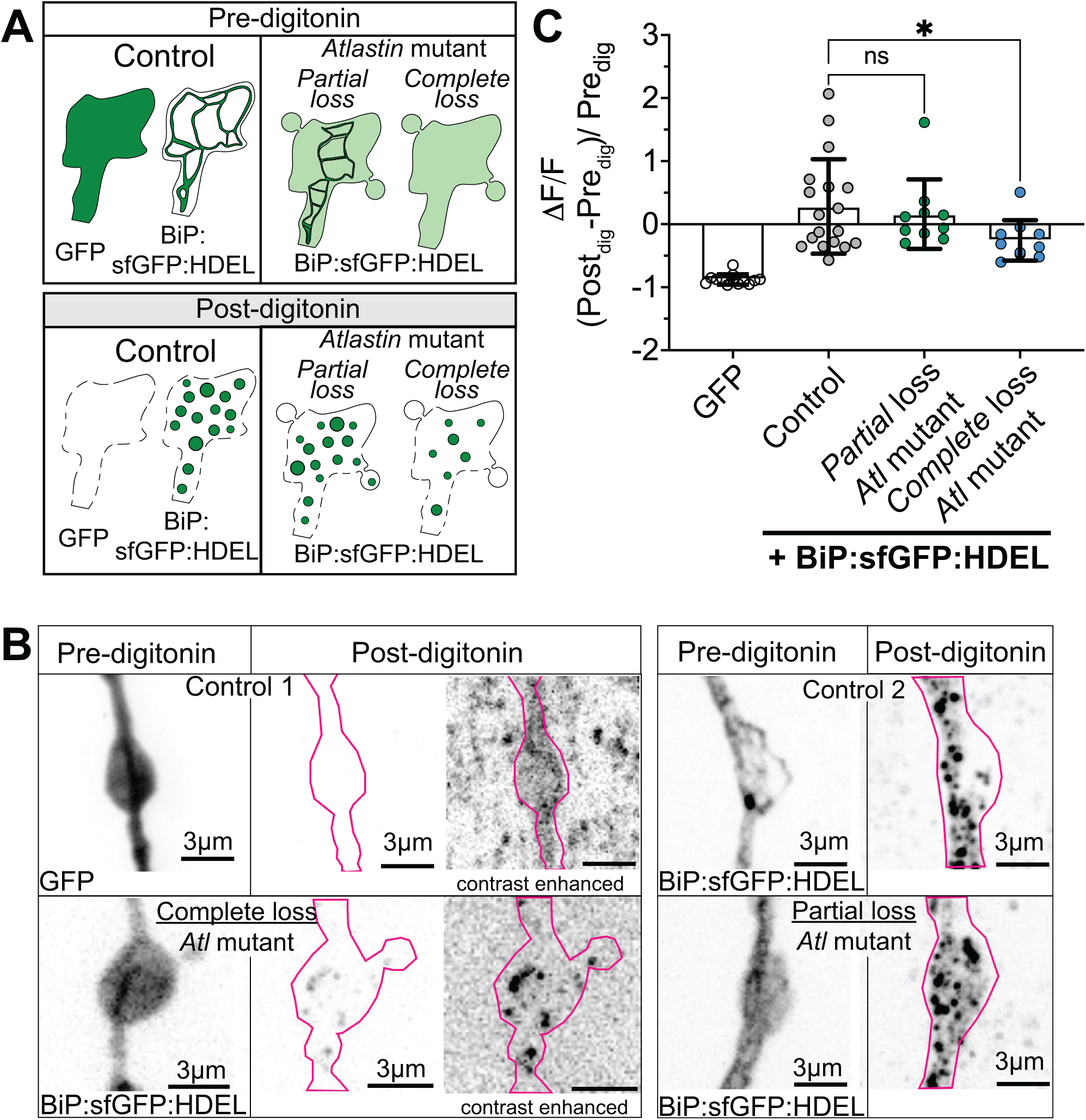
Luminal ER proteins displaced to the cytosol are sensitive to digitonin treatment. (**A**) Diagram of the expected distribution patterns before and after digitonin treatment of GFP and BiP:sfGFP:HDEL. Note that digitonin treatment causes ER fragmentation. (**B**) Maximum projections of Z-stacks acquired live with a spinning disk confocal of boutons expressing with GFP (control 1; highly sensitive to digitonin treatment), BiP:sfGFP:HDEL (control 2; resistant to digitonin treatment), BiP:sfGFP:HDEL in *Atlastin* mutants exhibiting partial loss and BiP:sfGFP:HDEL in *Atlastin* mutants exhibiting complete loss. (**C**) Fractional change of fluorescence intensity at presynaptic terminals expressing GFP (cytosolic protein) or BiP:sfGFP:HDEL in control and *Atlastin* mutants following digitonin treatment. Detailed genotype, replicate, and statistical information can be found in Supplementary Table 1.

We next performed a fluorescence recovery after photobleaching (FRAP) experiment to test the dynamics of BiP:sfGFP:HDEL in the presence of morphotrap_Int_. We photobleached *en passant* boutons using a defined region of 6.8 x 7.8 microns (dashed box in **Figure 8D**) to ensure that BiP:sfGFP:HDEL could recover from the ER networks surrounding the FRAP region (**Movies S20–S23**). We recorded movies of a single NMJ slice, imaging both markers. After photobleaching with a 405 nm laser, we captured recovery dynamics by imaging every 10 seconds for 2 minutes. We applied a one-association non-linear model to fit the recovery of BiP:sfGFP:HDEL or morphotrap_Int_ after FRAP, measuring the fraction of unbleached marker that reappeared in the FRAP region (mobile fraction). We found that morphotrap_Int_ itself recovered very slowly and plateaued at 19.6% in wild-type and 17.1% in *Atlastin* mutants (**Figure 8E**). We predicted that this slow diffusion of morphotrap_Int_ would restrict the movement of any cytosolic BiP:sfGFP:HDEL in *Atlastin* mutants. Indeed, we found that wild-type neurons expressing morphotrap_Int_ had a BiP:sfGFP:HDEL mobile fraction of 57.1%, while *Atlastin* mutant neurons had a smaller mobile fraction of 38.4% (**Figure 8F**). We hypothesize that the reduction in the mobile fraction in *Atlastin* mutants results from morphotrap_Int_ binding to BiP:sfGFP:HDEL in the cytosol. We next examined the recovery kinetics of BiP:sfGFP:HDEL in larvae co-expressing morphotrap_Int_ by measuring its fluorescence recovery half-life using the one-association non-linear model described previously (**Figure 8F**). We found that the recovery in controls was slower, with a half-life of 53.2 s, while *Atlastin* mutants had a faster recovery with a half-life of 45.8 s. The slightly faster half-life of the residual mobile BiP:sfGFP:HDEL in *Atlastin* mutants could reflect the presence of morphotrap_Int_ -free BiP:sfGFP:HDEL in the cytosol.

Finally, as another independent test of the subcellular localization of the ER lumenal marker, we examined whether BiP:sfGFP:HDEL in *Atlastin* mutants is sensitive to the mild detergent digitonin, which gently permeabilizes the plasma membrane and extracts proteins from the cytosol but not the ER (56). We found that digitonin treatment of *Atlastin* mutant presynaptic terminals with complete loss of BiP:sfGFP:HDEL resulted in a significant decrease in fluorescence intensity, further supporting the notion that the luminal ER marker is mislocalized to the cytosol in these synapses (**Figure 8—figure supplement 1**). Overall, our findings provide multiple lines of evidence that BiP:sfGFP:HDEL is displaced from the ER lumen to the cytosol in *Atlastin* mutants, potentially reflecting disruption to ER integrity and function at presynaptic terminals.

### *Atlastin* mutants have disrupted luminal ER protein dynamics

Having established that BiP:sfGFP:HDEL is displaced to the cytosol, we next performed FRAP of the marker alone to examine its cytosolic pool (which should recover rapidly relative to ER-localized proteins (57)) and the continuity of the ER network (continuous networks recover robustly while discontinuous ones recover poorly (57,58)). Using the same approach described above for morphotrap_Int_, we conducted FRAP analysis on single synaptic boutons co-expressing BiP:sfGFP:HDEL and RFP:mCD8 (which labels the neuronal plasma membrane). We applied a one-association non-linear model to fit the recovery of BiP:sfGFP:HDEL after FRAP to measure its mobile fraction and half-life, as previously described (**Movies S24–S27**). These measurements revealed distinct recovery patterns across genotypes. Partial loss *Atlastin* mutants showed both reduced mobile fractions (50.6%) and slower kinetics (half-life: 38.3 sec), while controls and complete loss mutants had similar, higher mobile fractions (65.7% and 66.3%) and faster recovery (26.9 sec and 14.6 sec, respectively) (**Figure 9A–B**). The reduced mobile fraction in partial loss *Atlastin* mutants likely reflects discontinuous ER networks (**Figure 9A (i-iv)**). The rapid recovery and absence of visible ER structures in complete loss *Atlastin* mutants suggests that BiP:sfGFP:HDEL is largely displaced into the cytosol. To determine whether observed changes are specific to ER proteins or reflect general protein mobility defects, we examined RFP:mCD8 as a negative control. Recovery could not be fitted to a non-linear equation because no plateau was observed (**Figure 9C**). Despite this limitation, RFP:mCD8 showed similar plasma membrane distribution and recovery rate across all genotypes, indicating no changes in plasma membrane organization. Overall, our findings reveal a novel pathogenic mechanism in which Atlastin dysfunction primarily affects ER protein distribution, potentially disrupting synaptic function through the mislocalization of essential ER proteins.

**Figure 9.**
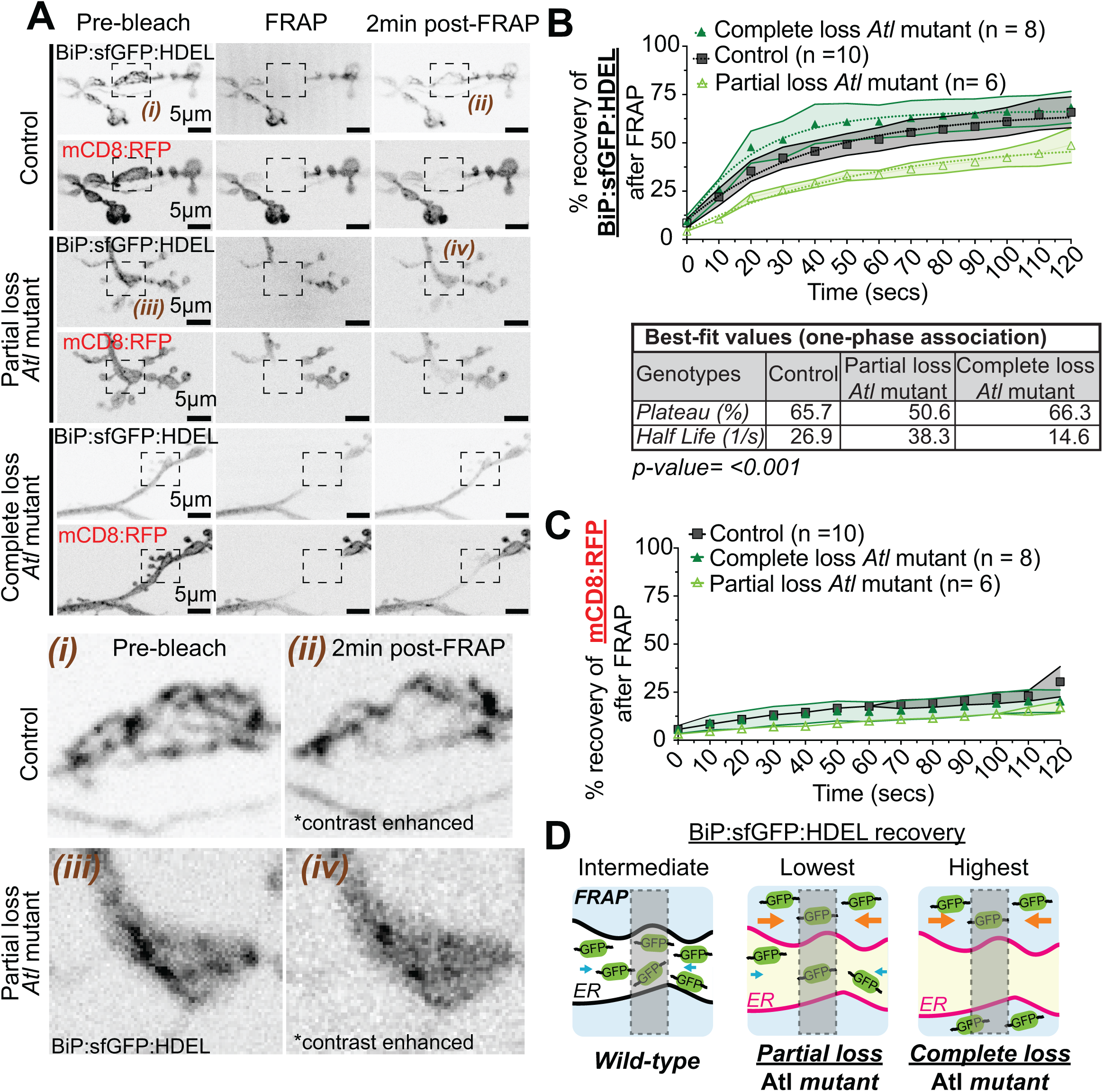
Atlastin mutants show disrupted luminal ER protein dynamics. (**A**) Representative slices from movies of controls and *Atlastin* mutants with partial loss or complete loss co-expressing mCD8:RFP and BiP:sfGFP:HDEL pre-bleach, immediately after FRAP, and 2 min post-FRAP (**Movies S24–27;** same dataset as Fig. 8B). The dashed box in (A) indicates areas that were photobleached and analyzed for recovery quantification in (B–C). (**B**) Mean recovery of BiP:sfGFP:HDEL in control (black squares) and *Atlastin* mutants with partial loss (empty lime green triangles) and complete loss (filled green triangles). Table shows non-linear fit one-phase association parameters. s.e.m. are represented by solid lines and shading: controls (n=10) have black lines with grey shading, *Atlastin* mutants with partial loss (n=6) have lime green lines and lime green shading, and *Atlastin* mutants with complete loss (n= 8) have green lines with light green shading. (**C)** Mean recovery of mCD8:RFP in control (black squares) and *Atlastin* mutants with partial loss (empty lime green triangles) and complete loss (filled green triangles). (**D**) Diagram of the recovery of BiP:sfGFP:HDEL in controls and *Atlastin* mutants with partial loss or complete loss. Blue arrows indicate diffusion of BiP:sfGFP:HDEL within the ER, while orange arrows indicate its diffusion in the cytosol. Arrow size correlates with rate of diffusion. Detailed genotype, replicate, and statistical information can be found in Supplementary Table 1.

### *Atlastin* mutants exhibit mild ER stress

Displacement of luminal proteins to the cytosol can be a downstream consequence of ER stress (59–62), prompting us to ask whether this pathway is activated in *Atlastin* mutants. We quantified endogenous BiP, an ER chaperone whose levels correlate with ER stress activation. We confirmed our anti-BiP antibody could detect ER stress by treating wild-type larvae with 5 to 50 mM DTT for 24 hours. This treatment significantly increased BiP protein levels in the brain at 50 mM DTT, but not those of another luminal ER protein, GP93 (**Figure 10A**). We were unable to perform this experiment on *Atlastin* mutant larvae because they did not eat the DTT-treated food, as evidenced by the absence of blue-dyed food in their gut compared to controls. Instead, we compared baseline endogenous BiP levels at NMJs in *Atlastin* mutants expressing no ER markers, BiP:sfGFP:HDEL, or tdTomato:Sec61β. Endogenous BiP had a punctate distribution compared to BiP:sfGFP:HDEL, likely reflecting its interaction with other proteins. *Atlastin* mutants showed increased BiP levels at the NMJ, suggesting a buildup of unfolded proteins and activation of ER stress (**Figure 10B–E**). Interestingly, overexpression of either BiP:sfGFP:HDEL or tdTomato:Sec61β blocked this increase in endogenous BiP levels, arguing against the possibility that overexpression of BiP:sfGFP:HDEL artificially induces ER stress. We conclude that Atlastin may have mild ER stress, which is not worsened by a higher translation load when transgenes are overexpressed.

**Figure 10.**
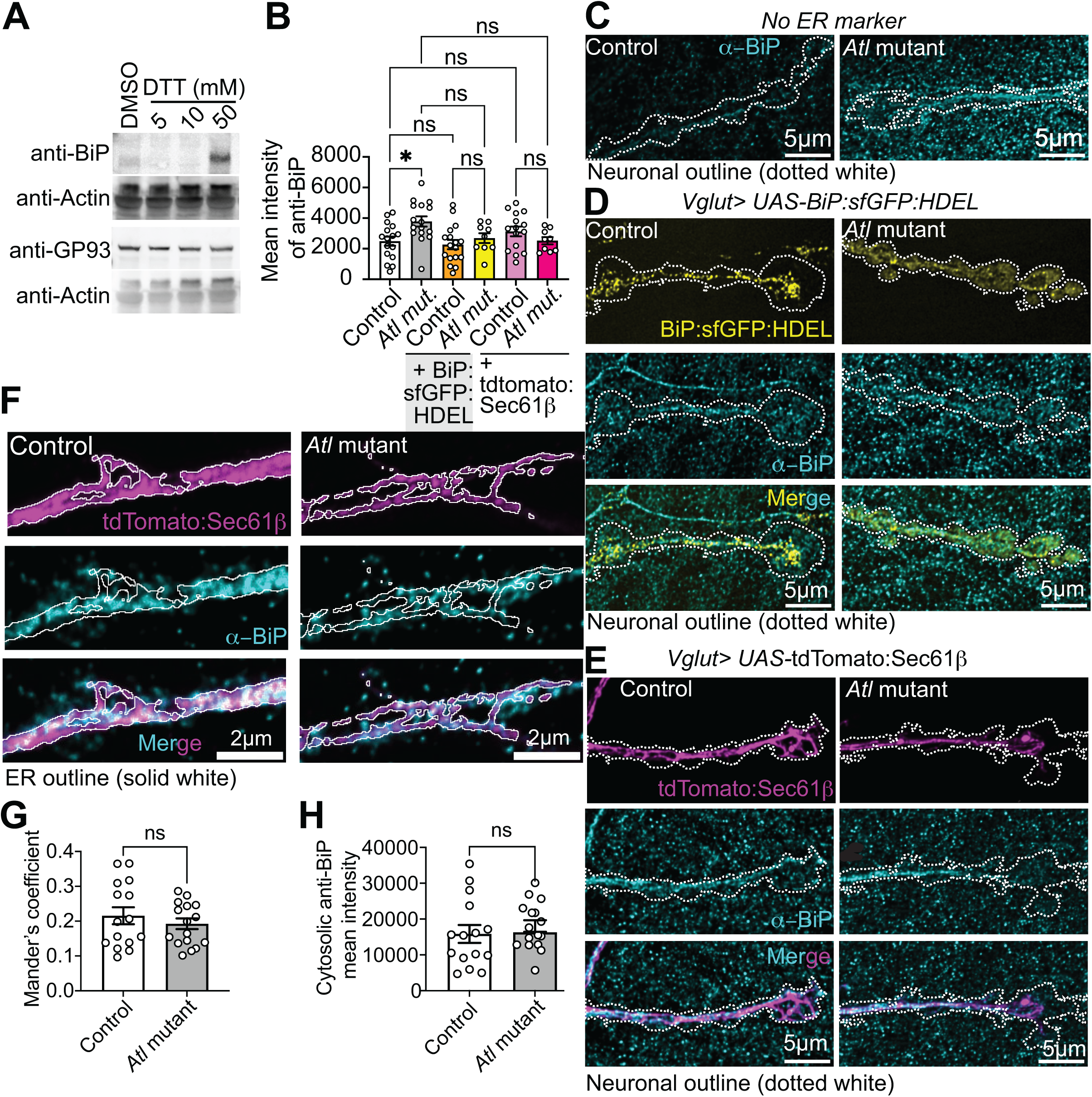
Endogenous BiP is elevated at synapses and is not displaced in *Atlastin* mutants. (**A**) Western blot following 24 hours of feeding wild-type animals with DMSO or DTT (5, 10, or 50 mM). (**B**) Mean intensity of α-BiP (magenta) in controls and *Atlastin* mutants with no ER marker, expressing BiP:sfGFP:HDEL or tdTomato:Sec61β. Representative SoRA maximum intensity projections of controls and *Atlastin* mutants with (**C**) no ER marker, (**D**) expressing BiP:sfGFP:HDEL, or (**E**) tdTomato:Sec61β. The dotted white line represents the neuronal outline. (**F**) Representative STED maximum intensity projections of controls and *Atlastin* mutants expressing tdTomato:Sec61β. The ER outline is represented by a solid white line. **(G)** Quantification of STED images via Mander’s correlation coefficient between anti-BiP and tdTomato:Sec61β and **(H)** Cytosolic levels of α-BiP in controls and *Atlastin* mutants. Detailed genotype, replicate, and statistical information can be found in Supplementary Table 1.

Next, we used the subcellular distribution of endogenous BiP to determine whether the displacement of the luminal ER marker BiP:sfGFP:HDEL reflects the displacement of all or only a subset of luminal ER proteins. We measured the Mander’s correlation coefficient of anti-BiP signal within tdTomato:Sec61β-positive regions in STED microscopy images of control and *Atlastin* mutants, and did not observe a significant difference (**Figure 10F–H**). These results indicate that only a subset of luminal ER proteins are displaced in *Atlastin* mutants.

### Proteasome inhibition increases ER-localized but not cytosolic BiP:sfGFP:HDEL

Proteins can be extruded from the ER lumen by the ER-Associated Degradation (ERAD) pathway, and are immediately targeted for ubiquitin-mediated degradation by the proteasome (63). We tested the hypothesis that BiP:sfGFP:HDEL accumulates in the cytosol of presynaptic terminals in *Atlastin* mutants due to defective proteasomal degradation. We found that ubiquitinated protein levels, as labeled by the FK1 antibody, were similar in control and *Atlastin* mutants, even when BiP:sfGFP:HDEL was expressed (**Figure 11A–B)**. To inhibit proteasomes, we treated larval fillets with 50 µM MG132 (or DMSO as a negative control) for 1 hour before fixation to allow the accumulation of ubiquitinated proteins destined for degradation. We labeled ubiquitinated proteins at fixed presynaptic terminals using the FK1 antibody and found comparably increased ubiquitination in both control and *Atlastin* mutant samples treated with MG132 (**Figure 11A–B**). Taken together, these results confirm the specificity of the anti-ubiquitin antibody in this preparation and suggest that proteasomes in *Atlastin* mutants are not broadly impaired.

**Figure 11.**
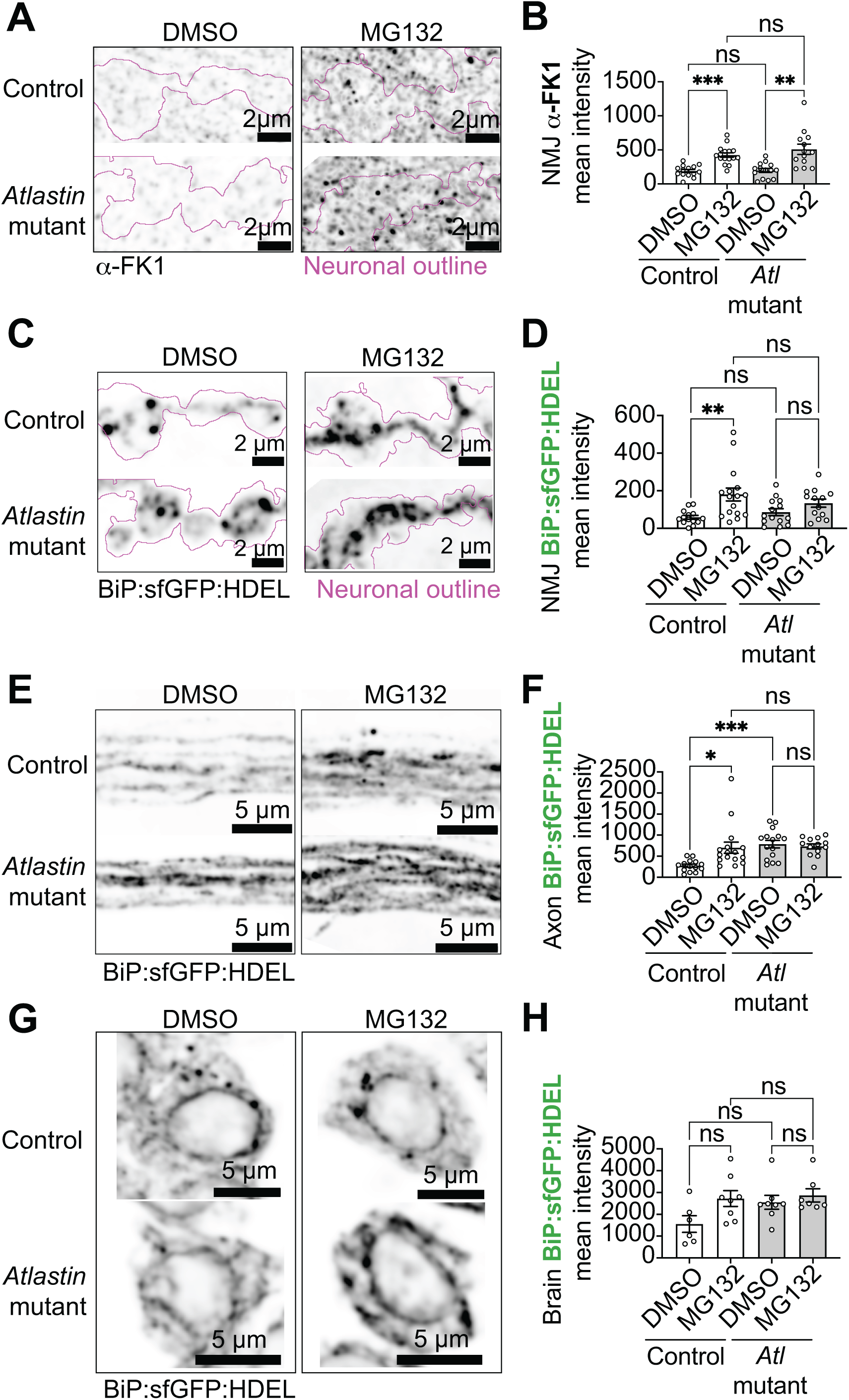
Cytosolic accumulation of BiP:sfGFP:HDEL is independent of proteasome activity. **(A)** SoRa Maximum intensity projections of α−FK1 immunostaining in control (top) and *Atlastin* mutant (bottom) NMJs treated with DMSO (left) or MG132 (right). **(B)** Quantification of α−FK1 mean intensity at NMJs in controls and *Atlastin* mutants treated with DMSO or MG132**. (C-G)** SoRa maximum intensity projections of BiP:sfGFP:HDEL in NMJs (**C**), axons (**E**) and cell bodies (**G**) in control (top) and *Atlastin* mutant (bottom) larvae treated with DMSO (left) or MG132 (right). Quantification of BiP:sfGFP:HDEL in NMJs (**D**), axons (**F**) and cell bodies (**H**). The data in this figure represents one dataset.

We next directly examined BiP:sfGFP:HDEL levels in these samples. BiP:sfGFP:HDEL accumulated in presynaptic terminals and axons of control animals following MG132 treatment but notably did not significantly accumulate in *Atlastin* mutants (**Figure 11C–F**). However, in contrast to presynaptic terminals, BiP:sfGFP:HDEL levels in axons of *Atlastin* mutants were already significantly elevated without MG132 treatment, suggesting baseline impairment of BiP:sfGFP:HDEL’s degradation. This is similar to our previous observation that tdTomato:Sec61β accumulates in *Atlastin* mutant axons (**Figure 4**). Finally, BiP:sfGFP:HDEL did not accumulate in the cell bodies of either controls or *Atlastin* mutants after MG132 treatment (**Figure 11G–H**). These results suggest that BiP:sfGFP:HDEL is more susceptible to degradation at axons and presynaptic terminals of controls compared to *Atlastin* mutants. Most importantly, MG132 treatment did not induce diffuse cytosolic BiP:sfGFP:HDEL in any compartment of control animals, including those that accumulated more BiP:sfGFP:HDEL following treatment. Instead, increased BiP:sfGFP:HDEL intensity observed in controls remained associated with intact ER networks rather than the cytosol. These results suggest subtle differences in proteostasis of BiP:sfGFP:HDEL in *Atlastin* mutants, but indicate that these defects are unlikely to contribute to the cytosolic displacement phenotype.

## Discussion

Here we reveal the architecture and dynamic behaviors of *Drosophila* neuronal ER networks through super-resolution *in vivo* imaging. This approach can be adapted to visualize fluorescently labeled proteins at high resolution in any larval tissue. Using this approach, we provided the first (to our knowledge) characterization of presynaptic ER network dynamics *in vivo*, establishing a foundation for investigating the functional significance of ER structural rearrangements. Our analysis of *Atlastin* mutants uncovered two unexpected phenotypes: First, these mutants maintain largely intact ER networks with only minor defects in dynamics; Second, they exhibit displacement of a luminal ER protein to the cytosol of presynaptic terminals but not in cell bodies or axons. These findings reveal that Atlastin serves distinct, synapse-specific functions in maintaining both ER structure and function, providing mechanistic insight into the functional deficits previously observed at *Atlastin* mutant synapses (21).

### ER network dynamics at presynaptic terminals

We observed three categories of ER dynamics at presynaptic terminals: (1) budding and fusion of vesicular structures, (2) ER tubule dynamics, and (3) dynamics of foci that are retained within the ER network. Future work is needed to examine the specific functions of these ER dynamics at presynaptic terminals, but below we outline the cell-biological processes that we speculate each category of dynamics may reflect. ER-derived vesicles could act as localized calcium signaling hubs (64) or serve as sites for compartmentalized local translation if they carry ribosomes (35,65). However, these vesicle-like structures are unlikely to be involved in exit from the ER via the secretory pathway or in ER-phagy (66) because we observe them re-fusing with the ER. These ER-derived vesicles are likely to involve ReepA and ReepB, the *Drosophila* orthologs of mammalian REEP1-4, which regulate ER vesicle formation in mammalian cells (67). Notably, while overexpression of Atlastin can regulate REEP vesicle fusion in mammalian systems (67), it is not essential for vesicle formation, suggesting similar regulatory relationships may exist between Atlastin and Reep genes in *Drosophila*. ER tubule dynamics may play multiple roles, such as maintaining the tension of the ER network (25,28) and forming ER-organelle contacts (29–31) that occur at the tips and edges of ER tubules and have diverse functions, including promoting mitochondrial and endosomal fission. To understand how these functions are regulated at presynaptic terminals, future studies could examine ER tubule interactions with other organelles using co-localization experiments or ER-organelle contact sensors (68). Foci dynamics within ER networks likely reflect changes in the dynamics of protein clusters at the ER membrane or within the lumen; foci dynamics occur at higher levels with the luminal ER marker than with the ER membrane marker, suggesting distinct organizational principles for luminal versus membrane proteins in presynaptic ER. We note that *Atlastin* mutants showed only mild ER dynamics defects, suggesting Atlastin plays a limited role in regulating presynaptic ER dynamic capacity.

### The presynaptic ER in *Atlastin* mutants forms architecturally normal but functionally discontinuous networks

A previous study reported that a luminal ER marker, BiP:sfGFP:HDEL, is diffusely distributed at *Atlastin* mutant presynaptic terminals, and interpreted this as extensive fragmentation of the ER network (21). However, this was not validated with an ER membrane marker or by electron microscopy, leaving it unclear whether the diffuse signal reflects a fragmented network or a redistribution of the marker itself. Using the membrane marker tdTomato:Sec61β, we found that the ER forms robust, well-organized networks at *Atlastin* mutant presynaptic terminals, with only mild defects in structure and dynamics. Further, we found that a substantial fraction of overexpressed BiP:sfGFP:HDEL is displaced from the ER lumen to the cytosol at *Atlastin* mutant synapses, using multiple independent approaches: capture of the marker by a cytosol-facing, plasma membrane-tethered nanobody, sensitivity to digitonin permeabilization, and rapid FRAP recovery consistent with free cytosolic diffusion.

Since the luminal and membrane markers report different structures, we considered which is more likely to reflect ER architecture in *Atlastin* mutants in the absence of any marker. We favor the hypothesis that ER networks in *Atlastin* mutants are similar to the tdTomato:Sec61β distribution, for the following reasons: First, tdTomato:Sec61β-expressing animals still show Atlastin-associated functional defects such as defective synaptic growth, indicating that Atlastin-dependent ER functions are disrupted without dramatic defects in ER structure. Second, the BiP:sfGFP:HDEL phenotype becomes progressively more severe throughout larval development, suggesting this marker’s displacement is a secondary effect rather than a direct reflection of ER shaping protein function. Third, the absence of BiP:sfGFP:HDEL displacement in *Reticulon 1* mutants, another key ER-shaping protein, strongly suggests that this phenotype is a specific consequence of Atlastin dysfunction rather than a general outcome of disrupting ER-shaping factors.

Although the ER networks in *Atlastin* mutants are architecturally normal, our FRAP data indicate that they are not fully continuous. This discontinuity could arise from various factors, such as protein blockages in the ER lumen, pinching of ER tubules, or other structural alterations that disrupt the continuity of the ER network without causing extensive fragmentation. By contrast, overall disruption of ER integrity seems unlikely to explain these observations: Catastrophic membrane failure would be incompatible with neuronal survival (69), and *Atlastin* mutant neurons show no evidence of apoptosis-induced ER permeability or the characteristic debris observed after neuronal death at the NMJ (70–72). Instead, we propose that changes in the ER network may lead to defects in the localization and function of ER-resident proteins. This hypothesis is supported by our observation that the levels of the ER membrane protein Sec61β are significantly reduced in *Atlastin* mutants.

Our work suggests that only a subset of luminal ER proteins may be displaced to the cytosol in *Atlastin* mutants: while we observed the displacement of the luminal marker BiP:sfGFP:HDEL, we were unable to identify an endogenous luminal ER protein exhibiting similar behavior. Initially, we hypothesized that endogenous BiP could be a good candidate for displacement because it is involved in ER stress and resides in the ER lumen. However, we did not observe displacement of endogenous BiP, suggesting that specific properties (such as ER retention and/or interactions with binding partners) determine the susceptibility of luminal proteins to cytosolic displacement.

### Mechanisms for displacement of a luminal ER protein to the cytosol

Our data shed light on how luminal proteins come to be found in the cytosol in *Atlastin* mutants. Several observations argue against the hypothesis that BiP:sfGFP:HDEL is aberrantly generated in the cytosol (rather than displaced from the ER to the cytosol) in *Atlastin* mutants: First, we do not observe cytosolic BiP:sfGFP:HDEL in cell bodies, where most protein translation is thought to occur. Second, we did not observe an increase in cytosolic BiP:sfGFP:HDEL in either controls or *Atlastin* mutants in our proteasome inhibition experiments, further suggesting that the protein is not generated in the cytosol. Third, the progressive nature of the phenotype over development (from 1^st^ to 3^rd^ instar) is consistent with the idea of protein displacement from the ER to the cytosol over time.

By what route might a luminal protein exit the ER? The accumulation of the endogenous ER chaperone BiP at *Atlastin* mutant presynaptic terminals is consistent with activation of ER stress responses such as the Unfolded Protein Response (UPR) or ER-Associated Degradation (ERAD) (59–61,73–75), and agrees with previous work showing UPR activation in *Atlastin* and other HSP-linked mutants (40,76,77). Importantly, however, the displaced BiP:sfGFP:HDEL retains fluorescence, indicating that the protein remains in a functional, folded state rather than being degraded. This aligns more closely with ER-to-cytosol signaling (ERCYS), in which luminal proteins are displaced into the cytosol as functional proteins capable of acquiring new cytosolic functions (61,62), rather than being degraded through the traditional ERAD pathway. Mechanistically, this is consistent with a recent study showing that the ATF6 and IRE1 branches of the UPR regulate ERCYS initiation (78), raising the possibility that the UPR activation we observe at *Atlastin* mutant terminals promotes ERCYS rather than ERAD. Importantly, ERCYS has only been demonstrated to date in yeast and in cancer cells, and our study raises the possibility that it also occurs in neurons *in vivo*.

Given these observations, what might trigger an ER stress response in *Atlastin* mutants? We hypothesize that reduced diffusion of BiP:sfGFP:HDEL in *Atlastin* mutants reflects broader changes in luminal dynamics, which might lead to altered calcium availability in presynaptic ER networks. This is consistent with studies in HeLa cells showing that ATL2/3 knockouts have reduced luminal content transport (33). Interestingly, ATL2/3 KOs had normal calcium storage levels but exhibited reduced local calcium release in response to IP_3_ uncaging, suggesting a deficit in local calcium dynamics. Whether impaired luminal diffusion, calcium dysregulation, or a combination of both is sufficient to initiate ER stress at synaptic terminals remains to be established.

### Synapse-specific vulnerability of the ER and its relevance to HSP

A striking feature of the displacement phenotype is its confinement to presynaptic terminals: the luminal marker is displaced to the cytosol at synapses while remaining correctly localized in cell bodies, axons, and body-wall muscles. However, synapse-specificity of the *Atlastin* mutant phenotype is not a unique property of the BiP:sfGFP:HDEL reporter. We observed a similarly local phenotype with the membrane marker tdTomato:Sec61β, which is reduced specifically at presynaptic terminals and not in cell bodies. Several features may render synaptic ER selectively vulnerable to loss of Atlastin function: Presynaptic ER is composed of thin, densely packed tubules, which may be especially dependent on Atlastin-mediated fusion to maintain continuity and proper distribution of ER membrane proteins. Synaptic ER is also physically remote from the cell body, where the bulk of protein synthesis and quality-control machinery resides, potentially limiting the terminal’s capacity to buffer local failures in luminal protein retention. We propose that synaptic ER normally operates close to a functional limit that is sensitive to loss of Atlastin, whereas better-buffered compartments such as the cell body are spared.

This synapse-specific vulnerability may be directly relevant to the pathogenesis of HSP, in which mutations in Atlastin cause one of the most common autosomal-dominant forms of the disease (15). HSP is a length-dependent axonopathy in which the longest neurons of the corticospinal tract degenerate progressively (17,52–54). We found that luminal ER protein retention fails specifically at presynaptic terminals, which are the compartments most distal from the cell body. This offers a cellular mechanism for the distal, progressive vulnerability characteristic of HSP, in which disruption of ER luminal function at presynaptic terminals, rather than gross ER structural collapse, may contribute to pathology. More broadly, our results underscore the importance of examining ER-shaping proteins in neurons, particularly at synapses, since the phenotypes we observe are not found in non-neuronal cells and non-synaptic compartments, where these proteins have most often been studied.

### Limitations of the study

A central limitation of this study is that the displacement phenotype was observed with an overexpressed reporter, raising the possibility that it reflects a property of the marker rather than of endogenous luminal ER proteins. Several observations argue against a simple artifact. First, it is not a generic consequence of overexpression stress: overexpressing either BiP:sfGFP:HDEL or tdTomato:Sec61β blocked rather than enhanced endogenous-BiP increase (**Fig. 10**), suggesting that overexpression is not directly correlated with ER stress. Second, it is not a generic feature of ER-shaping mutants: the same reporter localizes robustly to ER networks in *Reticulon1* mutants. Third, it is not likely a reporter-folding or aggregation defect: the marker localizes correctly in wild-type synapses, and displacement is *Atlastin*-specific and progressive. Confirming displacement of an endogenous luminal protein is an important goal for future work. However, several technical limitations will need to be overcome to achieve this goal. There are few tagged transgenes available for *Drosophila* luminal ER proteins. Detecting endogenous proteins with antibodies requires fixation and permeabilization, which notoriously disrupts ER structure and causes our reporter BiP:sfGFP:HDEL to collapse from a smooth distribution, as visualized by live imaging and FRAP, to a punctate distribution (compare **Fig. 6** to **Fig. 10**). Further, using antibodies rather than neuronally restricted transgenes makes it challenging to determine whether the signal originates from the neuron or from dense ER structures in the surrounding muscle. Some ER luminal proteins can be displaced by as little as 30%, and the sensitivity of our imaging assays may limit our ability to detect these small changes (61). By contrast, quantitative biochemical approaches such as fractionation, are not possible in our complex *in vivo* sample because neuronal ER proteins would mix with ER from other tissues upon homogenization. These limitations highlight the need for future studies to develop new tools and techniques to study the selective displacement of luminal ER proteins in *Atlastin* mutants.

## Methods

### Drosophila stocks

*Drosophila melanogaster* was cultured on a standard medium at 22–25°C. Detailed genotypes and sexes used for experiments are described in **Table S1**. GAL4 drivers used in this study include: Elav^C155^ (FlyBaseID: FBti0002575), Vglut-GAL4 (Bloomington *Drosophila* Stock Center (BDSC_24635), and BG57 (gift from Vivian Budnik). Other stocks used in this study are *atl*^2^ (gift from James McNew), *Reticulon1^18^*(gift from Cahir O’Kane), third chromosome deficiency (BDSC_7948), UAS-BiP:sfGFP:HDEL (BDSC_64748, BDSC_64749), 20XUAS-tdTomato:Sec61β (BDSC_64746), UAS-CD8:RFP (BDSC_27391), UAS-Atl RNAi (BDSC_36736), UAS-mCherry RNAi (BDSC_35785) and UAS-morphotrap_Int_ (gift from Chi-Kuang Yao).

### Construction of *in vivo* imaging slide for *Drosophila* larvae

To make the reusable imaging slide, Press-To-Seal Silicone Isolators (Grace Bio-labs; CQS-13R-2.0) were mounted onto rectangular glass slides, which were then filled from their centers to their tops with Krayden Dow Sylgard 184 Silicone (Thermo Fisher Scientific). Each slide with a Press-To-Seal Silicone Isolator was leveled to ensure the silicone cured evenly at room temperature. Minutien pins (Fine Science Tools) were cut with nail clippers and used to stretch the larvae and fix it in place. Pins were pressed into the cured Sylgard 184 Silicone, and ∼200 µL of hemolymph-like solution HL3.1 (44) were added. Next, a coverslip was placed on top of the dissected larvae, each corner was gently pressed down, and excess HL3.1 was removed with a Kimwipe, thus sealing the coverslip to the silicone isolator.

### Live imaging of luminal and membrane ER markers

1^st^ and wandering 3^rd^ instar larvae were dissected in Ca^2+^-free HL3.1 and axons were severed from the central nervous system. Dissections were performed in the live imaging slide described previously. Larvae were imaged at room temperature with 63X (n.a. 1.4) oil immersion objective on one of two microscopes, as indicated in Figure Legends: (1) an Airyscan LSM 880 microscope in super-resolution mode and Zen Black software. Raw image stacks were processed in Zen Blue or Zen Black using 3D Airyscan processing with automatic settings. (2) a Nikon super-resolution spinning disk SoRA system in W1-spinning disk mode, equipped with an Apo 60x Oil DIC N2 objective and set to a 12-bit depth. Images were collected using Nikon Elements AR software and deconvolved with Huygens deconvolution software.

Live fillets of 1^st^ and 3^rd^ instar larvae were stained with 1:250 dilution of 1.5 mg/mL α−HRP Alexa Fluor 647 (Jackson ImmunoResearch) in HL3.1 for 5 min followed by a 1 min wash. For 3^rd^ instar larvae, Z-stacks were collected of cell body clusters, axons, and presynaptic type Ib terminals at muscle 4 in segment A5 (unless otherwise noted). These data were used to measure the levels of luminal ER markers, to categorize ER network phenotypes, or for morphological measurements. For 1^st^ instar larvae, Z-stacks of presynaptic terminals at muscle 4 in segments A3-A5 were acquired and used to categorize ER network phenotypes.

To record the dynamics of the luminal ER marker BiP:sfGFP:HDEL, 16-bit Z-stacks of a single bouton (8 slices with 0.5 micron spacing) were collected for 100 frames at a frame rate of 0.73 seconds/stack. To record ER dynamics with the membrane ER marker tdTomato:Sec61β, 8-bit Z-stacks of a single bouton (20 slices with 0.1850 micron spacing) were collected for 60 frames at a frame rate of 0.92 seconds/stack.

### Fixed tissue staining

Wandering 3rd instar larvae were dissected in Ca²⁺-free HL3.1 and fixed for 17 min in 4% PFA. To prevent ER fragmentation, the fixative was prepared by diluting 16% PFA (EMS, SKU: 15710) with a 2X HL3.1 stock (140 mM NaCl, 10 mM KCl, 8 mM MgCl₂, 20 mM NaHCO₃, 10 mM trehalose, 230 mM sucrose, and 10 mM HEPES) and ddH₂O to a final concentration of 1X HL3.1 and 4% PFA, rather than diluting in water alone. Larvae were blocked and permeabilized overnight in PBS containing 0.25% Saponin, 2.5% normal goat serum (NGS), 2.5% bovine serum albumin (BSA), and 0.1% sodium azide. Fixed larvae were stained with primary antibodies at 4°C for 24 hrs and with 1:100 α− HRP Alexa Fluor 647 at room temperature for 2 hrs. Stained larvae were mounted in ProLong Diamond Antifade Mountant (#P36970; Thermo-Fisher Scientific, Waltham, MA, USA). Primary antibodies include: 1:500 FluoTag®-X4 α−RFP (#N0404, Nanotag Biotechnologies), 1:500 FluoTag®-X4 α−GFP (#N0304, Nanotag Biotechnologies), α−FK1 (#04-262, Sigma), Living Colors® α−mCherry Monoclonal Antibody (Takara; 632543) and 1:500 α−BiP (gift from Christopher Nicchitta).

Z-stacks were collected of *Drosophila* larval NMJs in muscle 4 of segment A5 at room temperature. One of three microscopes specified in the figure legend was used: (1) A Nikon Ni-E upright microscope equipped with 100x (n.a. 1.45) oil immersion objective, a Yokogawa CSU-W1 spinning-disk head, and an Andor iXon 897U EMCCD camera. (2) A Nikon super-resolution spinning disk SoRA system in W1-spinning disk mode, equipped with an Apo 60x Oil DIC N2 objective and set to a 12-bit depth. Images were collected using Nikon Elements AR software. (3) An Abberior 3D STED using a 60X silicone objective, mounted on an Olympus IX83 microscope and equipped with 405, 485, 561 and 640nm excitation lasers and a 775nm depletion laser.

### MG132 experiment

Larval fillets were incubated at room temperature in Schneider’s medium containing either DMSO vehicle control or 50 μM MG132 (diluted from 10 mM stock) for 1 hour at room temperature. Following treatment, samples were fixed, as previously described, and their cell bodies, axons, and NMJs were imaged.

### FRAP

Imaging was conducted using a Nikon super-resolution spinning disk SoRA system in W1-spinning disk mode, equipped with an Apo 60x Oil DIC N2 objective and set to a 12-bit depth. We photobleached boutons in the middle of a synaptic branch to ensure BiP:sfGFP:HDEL could recover from both the top and bottom of the photobleached region. To capture the rapid recovery of BiP:sfGFP:HDEL, we imaged a single slice of NMJs at muscle 4 and sequentially acquired 488 and 561 channels with an exposure time of 100ms. For all NMJs, we photobleached single boutons within a 6.8x7.8 microns region in the middle of neuronal branch. Each movie acquired used an ND sequence acquisition consisting of three phases: (1) we acquired five frames at maximum speed (100Mhz) before photobleaching; (2) we photobleached BiP:sfGFP:HDEL and RFP:mCD8 or morphotrap_Int_ using a 405 nm depletion laser with a dwell time of 100µs for a duration of 2s; and (3) we sequentially imaged the recovery of BiP:sfGFP:HDEL and RFP:mCD8 or morphotrap_Int_ with a 10s interval for 2 minutes for experiments in **Figure 8** and 5 minutes for experiments in **Figure 9**.

### Digitonin treatment

Digitonin experiments were conducted in live fillets from larvae expressing the luminal ER marker, UAS-BiP:sfGFP:HDEL, under the control of the neuronal driver Vglut-GAL4. Fillets were stained with 1:500 1.5 mg/ml α−HRP Alexa Fluor 594 (Jackson ImmunoResearch) or 1:250 α−HRP Alexa Fluor 647 in HL3.1 for 5 min, followed by a 1 min HL3.1 wash. Next, pre-digitonin Z-stacks were acquired from muscle 4 NMJs from segments A3 to A5 (both hemisegments) using the spinning disk confocal microscope described above. Fillets were subsequently incubated with 0.05% digitonin in HL3.1 for 5 min, then rinsed with HL3.1 and re-incubated with α−HRP Alexa Fluor 594 or α−HRP Alexa Fluor 647 for 5 min. Samples were then washed for 1 min with HL3.1. Finally, post-digitonin Z-stacks were acquired of the same NMJs.

### Image analysis

All processing of images and videos was performed using ImageJ/Fiji. Debris and axon bundles were manually removed from all images before analysis of maximum projections or Z-stacks. Neuronal masks were created by first conducting a background subtraction of the image (rolling ball, 50 pixel radius) and then introducing a gaussian blur before thresholding using Otsu or Li algorithms. <u>For live image analysis,</u> maximum intensity projections of each time point were corrected for (1) photobleaching using histogram matching and (2) sample drift or muscle contraction using the StackReg plugin (79).

1. *Categorization of ER dynamics:* For dynamics analysis, we used the same dataset as the control in **Figure 2**, **Figure 5**. Maximum intensity projection timelapse movies were blinded and replayed 3-4 times before categorization. We identified three distinct categories of ER dynamics in presynaptic terminals of *Drosophila* motor neurons (note that a single bouton can exhibit multiple types of dynamics): (1) **<u>ER budding</u>** – small vesicle-like structures emerging from the ER network and re-fusing in a different region; (2) **<u>ER tubule dynamics</u>**, encompassing four subcategories as follows: (a) <u>Tubule displacement</u>, involving tubules that changed their localization within boutons without detaching from the ER network; (b) <u>Tubule extension</u>, where an ER tubule emerged and extended from the ER network; (c) <u>Tubule retraction</u>, characterized by the retraction and disappearance of an ER tubule from the ER network; and (d) <u>Tubule</u> <u>extension and retraction</u>, indicating an ER tubule both extending and retracting from the ER network within the timelapse. (3) **<u>Foci dynamics</u>** – round, bright puncta moving within the ER network and (4) **<u>static</u>** – boutons that lacked observable dynamics.
2. *Analysis of NMJ morphology:* Satellite and total bouton numbers were manually counted from blinded maximum intensity projections of fixed and α−HRP labeled larvae. Satellite boutons were defined as any string of 5 or fewer boutons extending from the main axis of the NMJ (**Figure 3A**).
3. *Presynaptic levels of ER membrane* marker tdTomato:Sec61β (***Figure 3C–D*, *Figure 10E***): *A* custom FIJI macro consisting of the following steps was employed on maximum intensity projections,: (1) An NMJ mask was created using the α−HRP neuronal marker; (2) The integrated density of the ER membrane marker tdTomato:Sec61β was measured within the α−HRP mask; (3) The mean intensity of the ER membrane marker at presynaptic terminals was calculated by dividing the tdTomato:Sec61β integrated density by the area of the presynaptic terminal.
4. *Categorization of luminal ER marker distribution:* Images were blinded before analysis to ensure unbiased qualitative analysis of luminal ER marker BiP:sfGFP:HDEL in controls and *Atlastin* mutants. Maximum intensity projections were then classified into three phenotypic categories: (1) **<u>“Partial loss”</u>** - luminal ER marker localizes to ER networks but is partially displaced to the cytosol, (2) **<u>“Complete loss”</u>**- luminal ER marker mostly displaced to the cytosol and ER networks are faint or not detectable, and (3) **<u>“No phenotype”</u>**- luminal ER marker completely localizes to ER networks.
5. *Presynaptic levels of luminal ER proteins at* Drosophila *motor neurons:* A custom FIJI macro consisting of the following steps was employed on Z-stacks: (1) We used as a mask BiP:sfGFP:HDEL (**Figure 6D–E**) or α−HRP (**Figures 6F**, **7B–C, 7F–H**, **10C–E**). (2) Using either mask, we measured its volume and the integrated density of various markers (e.g., BiP:sfGFP:HDEL, α−FK1, α−BiP). (3) We calculated the mean intensity of the marker at the NMJ by dividing the integrated density by the volume of the presynaptic terminal.
6. *BiP:sfGFP:HDEL, RFP:mCD8 or morphotrap_Int_ levels before and after photobleaching (****Figure 8-9***): Images were blinded for analysis. The 20 frames corresponding to the photobleaching event were removed since this prevented the Stackreg plugin from working correctly. Movies with uncorrectable drift were excluded. For the analysis, we drew 3 ROIs: the photobleached region, a region away from the bleach site, and the background. A Fiji macro was then used to measure the mean intensities in these 3 regions for each movie. The mean intensities at the site of photobleaching of BiP:sfGFP:HDEL, RFP:mCD8 or morphotrap_Int_ were background subtracted and normalized to the mean intensity of the non-bleached region.
7. *Presynaptic levels of cytosolic or luminal ER proteins pre- and post-digitonin treatment (****Figure 8—figure supplement 1***): Images were blinded and maximum projections were created for further analysis. Neuronal outlines were obtained by manually tracing the α−HRP-positive region in FIJI. Next, the integrated density of GFP or BiP:sfGFP:HDEL was quantified within the manually traced neuronal outlines. Mean intensity was calculated by dividing the integrated density by the area of the presynaptic terminal.
8. *Measuring the subcellular localization of endogenous BiP (****Figure 10G***): We calculated the Mander’s correlation coefficient by measuring the raw integrated density of endogenous BiP within the tdTomato:Sec61β mask and dividing it by the raw integrated density of BiP within the HRP mask in controls and *Atlastin* mutants.
9. *Quantification of cytosolic α−BiP levels (****Figure 10H***): We subtracted the mean intensity of α−BiP in the tdTomato:Sec61β mask from that in the HRP mask.

### Statistics

GraphPad Prism software was used to perform statistical analyses and generate graphs. Statistical analyses were conducted using parametric tests (e.g. one-way ANOVA, Student’s t-test) for normally distributed data, and non-parametric tests (e.g. Kruskal-Wallis, Mann-Whitney tests) if the data was not normally distributed. Chi-squared tests were used for categorical data. p-values were set to *<0.05, **<0.01, ***<0.001.

### Western blot

Larvae were grown at 25°C in Formula 4-24® Instant *Drosophila* Medium (Blue) (Carolina Biological Supply), prepared following the manufacturer’s instructions and supplemented with DMSO or DTT (5, 10, or 50mM). Five 3^rd^ instar larvae brains were dissected in HL3.1 solution without calcium. 50µL of 1x Laemmli Sample Buffer (#1610737, Bio-Rad) was used to prepare protein samples for Western blot. Transfers were performed using the Trans-Blot® Turbo™ Transfer System (Bio-Rad). Nitrocellulose membranes were blocked for 30 min with Blocking Buffer for Fluorescent Western Blotting (MB-070, Rockland). Primary antibodies used were 1:1000 α−BiP (a gift from Christopher Nicchitta), 1:1000 α−GP93 (a gift from Christopher Nicchitta), and 1:1000 α−actin (JLA20, Developmental Studies Hybridoma Bank).

## Acknowledgements

We thank the Brandeis Light Microscopy Facility (RRID:SCR_025892) for their support and assistance in this work. We also thank the Bloomington *Drosophila* Stock Center (Indiana University, Bloomington, IN, NIH P40OD018537), Christopher Nicchitta, Cahir O’Kane and James McNew for fly lines, Steve Del Signore for help with image analysis, and Bruce Goode for use of the SoRA microscope. We also thank Chi-Kuang Yao for sharing reagents and stimulating discussion. This work was supported by NINDS grants R01 NS103967 to A.A.R., grant S10 OD034223 for the Abberior Facility Line STED microscope, and T32 NS007292 and K99 NS136720 to M.C.Q.F.

## Movie legends

**Movies S1–3:** Movies acquired at a frame rate of 0.73 frames per second of presynaptic terminals expressing the ER lumen marker BiP:sfGFP:HDEL. These movies are played at a frame rate of 10 frames per second and the scale bar is 1 µm.

**Movies S4–6.** Movies acquired at a frame rate of 0.65 frames per second of presynaptic terminals expressing the ER membrane marker tdTomato:Sec61β. These movies are played at a frame rate of 10 frames per second and the scale bar is 1 µm.

**Movie S7:** Example movie of ER budding at presynaptic terminals expressing the ER membrane marker tdTomato:Sec61β. This movie is played at 5 frames per second, and the scale bar is 0.5 µm.

**Movie S8:** Example movie of ER foci dynamics at presynaptic terminals expressing the ER membrane marker tdTomato:Sec61β. This movie is played at 5 frames per second, and the scale bar is 0.5 µm.

**Movie S9:** Example movie of ER displacement tubule dynamics at presynaptic terminals expressing the ER membrane marker tdTomato:Sec61β. This movie is played at 5 frames per second, and the scale bar is 0.5 µm.

**Movie S10:** Example movie of ER extension tubule dynamics at presynaptic terminals expressing the ER membrane marker tdTomato:Sec61β. This movie is played at 5 frames per second, and the scale bar is 0.5 µm.

**Movie S11:** Example movie of ER retraction and extension tubule dynamics at presynaptic terminals expressing the ER membrane marker tdTomato:Sec61β. This movie is played at 5 frames per second, and the scale bar is 0.5 µm.

**Movie S12:** Example movie of a static proximal bouton of controls expressing the ER membrane marker tdTomato:Sec61β acquired at a frame rate of 0.92 seconds/stack. The movie is played at a frame rate of 5 frames per second and the scale bar is 2 µm.

**Movie S13:** Example movie of a dynamic proximal bouton of controls expressing the ER membrane marker tdTomato:Sec61β acquired at a frame rate of 0.92 seconds/stack. The movie is played at a frame rate of 5 frames per second and the scale bar is 2 µm.

**Movie S14:** Example movie of a static terminal bouton of controls expressing the ER membrane marker tdTomato:Sec61β acquired at a frame rate of 0.92 seconds/stack. The movie is played at a frame rate of 5 frames per second and the scale bar is 2 µm.

**Movie S15:** Example movie of a dynamic terminal bouton of controls expressing the ER membrane marker tdTomato:Sec61β acquired at a frame rate of 0.92 seconds/stack. The movie is played at a frame rate of 5 frames per second and the scale bar is 2 µm.

**Movie S16:** Example movie of a static proximal bouton of *Atlastin* mutants expressing the ER membrane marker tdTomato:Sec61β acquired at a frame rate of 0.92 seconds/stack. The movie is played at a frame rate of 5 frames per second and the scale bar is 2 µm.

**Movie S17:** Example movie of a dynamic proximal bouton of *Atlastin* mutants expressing the ER membrane marker tdTomato:Sec61β acquired at a frame rate of 0.92 seconds/stack. The movie is played at a frame rate of 5 frames per second and the scale bar is 2 µm.

**Movie S18:** Example movie of a static terminal bouton of *Atlastin* mutants expressing the ER membrane marker tdTomato:Sec61β acquired at a frame rate of 0.92 seconds/stack. The movie is played at a frame rate of 5 frames per second and the scale bar is 2 µm.

**Movie S19:** Example movie of a dynamic terminal bouton of *Atlastin* mutants expressing the ER membrane marker tdTomato:Sec61β acquired at a frame rate of 0.92 seconds/stack. The movie is played at a frame rate of 5 frames per second and the scale bar is 2 µm.

**Movie S20:** Representative movie of a control NMJ expressing morphotrap_Int_ pre-bleach and after FRAP every 10 secs for 5 min. The movie is played at a frame rate of 5 frames per second, and the scale bar is 5 µm.

**Movie S21:** Representative movie of a control NMJ expressing BiP:sfGFP:HDEL pre-bleach and after FRAP every 10 secs for 5 min. The movie is played at a frame rate of 5 frames per second, and the scale bar is 5 µm.

**Movie S22:** Representative movie of a *Atlastin* mutant NMJ expressing morphotrap_Int_ pre-bleach and after FRAP every 10 secs for 5 min. The movie is played at a frame rate of 5 frames per second, and the scale bar is 5 µm.

**Movie S23:** Representative movie of a *Atlastin* mutant NMJ expressing BiP:sfGFP:HDEL pre-bleach and after FRAP every 10 secs for 5 min. The movie is played at a frame rate of 5 frames per second, and the scale bar is 5 µm.

**Movie S24:** Representative movie of an *Atlastin* mutant NMJ with partial loss expressing BiP:sfGFP:HDEL pre-bleach and after FRAP every 10 secs for 2 min. The movie is played at a frame rate of 5 frames per second, and the scale bar is 5 µm.

**Movie S25:** Representative movie of an *Atlastin* mutant NMJ with partial loss expressing mCD8:RFP pre-bleach and after FRAP every 10 secs for 2 min. The movie is played at a frame rate of 5 frames per second, and the scale bar is 5 µm.

**Movie S26:** Representative movie of an *Atlastin* mutant NMJ with complete loss expressing BiP:sfGFP:HDEL pre-bleach and after FRAP every 10 secs for 2 min. The movie is played at a frame rate of 5 frames per second, and the scale bar is 5 µm.

