## Supplementary Table 1 for "Localized disruption of the presynaptic endoplasmic reticulum in *Atlastin* mutants"

### Table S1 – Detailed genotypes and statistics reporting

Experiments were done at muscle 4 unless otherwise noted. All analyses were performed in Fiji.

| Figure/Panel | Genotype (Condition/Label) | N (# larvae, # boutons) | Measurement | # Datasets | Statistical Test(s) |
|---|---|---|---|---|---|
| **Figure 2B–D**<br>Dynamics of ER luminal and membrane markers | Vglut-GAL4;UAS-BiP:sfGFP:HDEL/+ (ER lumen)<br><br>Vglut-GAL4;;UAS-tdTomato:Sec61β/+ (ER membrane) | 2 larvae 10 boutons<br><br>4 larvae, 14 boutons | ER dynamics (qualitative) | 1 | N/A |
| **Figure 2E–G**<br>ER dynamics | Vglut-GAL4;;UAS-tdTomato:Sec61β/+ | 18 larvae<br>75 terminal boutons, 29 *en passant* boutons | ER dynamics | 1 | N/A |
| **Figure 3B**<br>Count of satellite boutons | Vglut-GAL4;UAS-tdTomato:Sec61β/+ (Control)<br><br>Vglut-GAL4;UAS-tdTomato:Sec61β/+;atl$^{2}$/atl$^{def}$ (*Atlastin* mutant) | 8 larvae, 30 NMJs<br><br>8 larvae, 30 NMJs | Count of satellite boutons | 1 | Unpaired Mann-Whitney test, P<0.0001 |
| **Figure 3C–D**<br>Overall levels of ER membrane markers at nerve terminals of *Atlastin* mutants | Vglut-GAL4;UAS-tdTomato:Sec61β/+ (Control)<br><br>Vglut-GAL4;UAS-tdTomato:Sec61β/+;atl$^{2}$/atl$^{def}$ (*Atlastin* mutant) | 8 larvae, 30 NMJs<br><br>8 larvae, 30 NMJs | (C) CoV<br>(D) Mean intensity of marker | 1 | (C) Unpaired Mann-Whitney test, P<0.0001<br><br>(D) Unpaired t-test, P<0.05 |
| **Figure 3F**<br>Overall levels of ER membrane markers at nerve terminals of *Reticulon1* mutants | Vglut-GAL4;;UAS-tdTomato:Sec61β/+ (Control)<br><br>Vglut-GAL4;Reticulon$^{18}$/Reticulon$^{18}$;UAS-tdTomato:Sec61β/+ (*Reticulon18* mutant) | 10 larvae, 19 NMJs<br><br>11 larvae, 18 NMJs | Mean intensity of marker | 1 | Unpaired Mann-Whitney test, P<0.0001 |
| **Figure 4C–H**<br>Overall levels of ER membrane markers at axon bundles and soma clusters in *Atlastin* mutants | Vglut-GAL4;UAS-tdTomato:Sec61β/+ (Control) | Cell bodies: 8 larvae, 26 clusters<br>Axon bundle: 6 larvae, 29 NMJs<br><br>Cell bodies: 5 larvae, 33 clusters | (C, F) Mean intensity of marker<br>(D, G) Area (μm²)<br>(E, H) CoV | 1 | (C-H) Unpaired Mann-Whitney test, P<0.0001 |

| Figure/Panel | Genotype (Condition/Label) | N (# larvae, # boutons) | Measurement | # Datasets | Statistical Test(s) |
|---|---|---|---|---|---|
| | Vglut-GAL4;UAS-tdTomato:Sec61β/+;$atl^2/atl^{def}$ (*Atlastin* mutant) | Axon bundle: 4 larvae, 32 NMJs | | | |
| **Figure 5B–F**<br>ER dynamics in *Atlastin* mutants | Vglut-GAL4;UAS-tdTomato:Sec61β/+ (Control)<br><br>Vglut-GAL4;UAS-tdTomato:Sec61β/+;$atl^2/atl^{def}$ (*Atlastin* mutant) | 18 larvae;<br>- 75 terminal boutons (67 dynamic, 8 static), -<br>- 29 en passant boutons (11 dynamic, 18 static)<br>(same dataset as Figure 2E–G)<br><br>20 larvae;<br>- 83 terminal boutons (56 dynamic, 27 static),<br>- 36 en passant boutons (10 dynamic, 26 static) | (B) Categories: dynamic or static, in terminal and en passant boutons<br><br>(C) ER dynamic types in terminal boutons (budding, foci dynamics, or ER tubule dynamics)<br><br>(D) ER dynamic types in en passant boutons (budding, foci dynamics, or ER tubule dynamics)<br><br>(E) ER tubule dynamics: displacement, extension, retraction, or both extension and retraction<br><br>(F) Number of dynamic categories per terminal bouton | 1 | Chi-square, P<0.0001 |
| **Figure 6C**<br>Categorization of ER network phenotypes in *Atlastin* mutants expressing luminal ER markers | Vglut-GAL4;UAS-BiP:sfGFP:HDEL/+ (Control)<br><br>Vglut-GAL4;UAS-BiP:sfGFP:HDEL/+;$atl^2/atl^{def}$ (*Atlastin* mutant) | 6 larvae, 20 NMJs<br><br>10 larvae, 23 NMJs | Categories: no phenotype, partial, or complete | 2 | Chi-square, P<0.0001 |
| **Figure 6D–F**<br>Overall levels of ER lumen markers at nerve terminals of *Atlastin* mutants | Vglut-GAL4;UAS-BiP:sfGFP:HDEL/+ (Control)<br><br>Vglut-GAL4;UAS-BiP:sfGFP:HDEL/+;$atl^2/atl^{def}$ (*Atlastin* mutant) | 6 larvae, 20 NMJs<br><br>10 larvae, 23 NMJs | (D, E) Mean intensity of marker<br>(F) CoV | 2 | (D, E) Unpaired Mann-Whitney test, P<0.0001<br>s<br>(F) Unpaired t-test, P<0.05 |

| Figure/Panel | Genotype (Condition/Label) | N (# larvae, # boutons) | Measurement | # Datasets | Statistical Test(s) |
|---|---|---|---|---|---|
| **Figure 6G**<br>Overall levels and CoV of ER lumen markers at nerve terminals of *Reticulon1* mutants | Vglut-GAL4;;UAS-BiP:sfGFP:HDEL/+ (Control)<br><br>Vglut-GAL4;Reticulon$^{18}$/Reticulon$^{18}$;UAS-BiP:sfGFP:HDEL/+ (*Reticulon18* mutant) | 4 larvae, 17 NMJs<br><br>6 larvae, 20 NMJs | Mean intensity of marker | 1 | Unpaired Mann-Whitney test, P<0.0001 |
| **Figure 7B–C**<br>Overall levels and CoV of ER lumen markers at nerve terminals of neuronal *Atlastin* knockdown | C155;UAS-AtlRNAi/+;UAS-BiP:sfGFP:HDEL/+ (*Atlastin* neuronal knockdown)<br><br>C155;;UAS-BiP:sfGFP:HDEL/UAS-mCherryRNAi (Control) | 6 larvae, 19 NMJs<br><br>4 larvae, 19 NMJs | (B) Mean intensity of marker<br>(C) CoV | 1 | (B) Unpaired Mann-Whitney test, P<0.0001<br><br>(C) Unpaired Mann-Whitney test, P<0.0001 |
| **Figure 7H**<br>Categorization of ER network phenotypes in 1st instar larvae, *Atlastin* mutants expressing a luminal ER marker | Vglut-GAL4;UAS-BiP:sfGFP:HDEL/+ (Control)<br><br>Vglut-GAL4;UAS-BiP:sfGFP:HDEL/+;atl$^{2}$/atl$^{def}$ (*Atlastin* mutant) | 7 larvae, 16 NMJs<br><br>6 larvae, 16 NMJs | Categories: no phenotype, partial, or complete | 1 | Chi-square, P<0.0001 |
| **Figure 8B**<br>Categorization of ER network phenotypes with RFP:mCD8 or Morphotrap$^{Int}$ co-expression | Vglut-GAL4;UAS-BiP:sfGFP:HDEL/UAS-RFP:mCD8 (Control, RFP:mCD8)<br><br>Vglut-GAL4;UAS-BiP:sfGFP:HDEL/UAS-RFP:mCD8;atl$^{2}$/atl$^{def}$ (*Atlastin* mutant, RFP:mCD8)<br><br>Vglut-GAL4;UAS-BiP:sfGFP:HDEL/UAS-Morphotrap$^{Int}$ (Control, Morphotrap$^{Int}$)<br><br>Vglut-GAL4;UAS-BiP:sfGFP:HDEL/UAS-Morphotrap$^{Int}$;atl$^{2}$/atl$^{def}$ (*Atlastin* mutant, Morphotrap$^{Int}$) | 4 larvae, 27 NMJs<br><br>4 larvae, 28 NMJs<br><br>3 larvae, 24 NMJs<br><br>6 larvae, 32 NMJs | Categories: no phenotype, partial, or complete | 1 | Chi-square, P<0.0001 |
| **Figure 8C**<br>Mean intensity of BiP:sfGFP:HDEL at nerve terminals co- | Vglut-GAL4;UAS-BiP:sfGFP:HDEL/UAS-RFP:mCD8 (Control, RFP:mCD8) | 4 larvae, 27 NMJs<br><br>4 larvae, 28 NMJs | Mean intensity of marker | 1 | Unpaired; one-way ANOVA (Kruskal-Wallis test), P<0.05 |

| Figure/Panel | Genotype (Condition/Label) | N (# larvae, # boutons) | Measurement | # Datasets | Statistical Test(s) |
|---|---|---|---|---|---|
| expressing RFP:mCD8 or Morphotrap$^{Int}$, in controls or *Atlastin* mutants | Vglut-GAL4;UAS-BiP:sfGFP:HDEL/UAS-RFP:mCD8;atl$^{2}$/atl$^{def}$ (*Atlastin* mutant, RFP:mCD8)<br><br>Vglut-GAL4;UAS-BiP:sfGFP:HDEL/UAS-Morphotrap$^{Int}$ (Control, Morphotrap$^{Int}$)<br><br>Vglut-GAL4;UAS-BiP:sfGFP:HDEL/UAS-Morphotrap$^{Int}$;atl$^{2}$/atl$^{def}$ (*Atlastin* mutant, Morphotrap$^{Int}$) | 3 larvae, 24 NMJs<br><br>6 larvae, 32 NMJs | | | |
| **Figure 8E–F** FRAP analysis of BiP:sfGFP:HDEL and Morphotrap$^{Int}$ in controls and *Atl* mutants | Vglut-GAL4/+;UAS-BiP:sfGFP:HDEL/UAS-Morphotrap$^{Int}$ (Control)<br><br>Vglut-GAL4/+;UAS-BiP:sfGFP:HDEL/UAS-Morphotrap$^{Int}$;atl$^{2}$/atl$^{def}$ (*Atlastin* mutant) | 4 larvae, 16 NMJs<br><br>6 larvae, 19 NMJs | Non-linear fit (one-phase association equation) | 1 | Extra sum-of-squares F test |
| **Figure 8_Suppl. C** Mean intensity of luminal ER markers in control and *Atlastin* mutants, post-digitonin treatment | Vglut-GAL4/+;UAS-GFP/+ (Control 1)<br><br>Vglut-GAL4/+;UAS-BiP:sfGFP:HDEL/+ (Control 2)<br><br>Vglut-GAL4/+;UAS-BiP:sfGFP:HDEL/+;atl$^{2}$/atl$^{def}$ (*Atlastin* mutant) | 4 larvae, 13 NMJs<br><br>5 larvae, 18 NMJs<br><br>5 larvae, 20 NMJs | Change in fluorescence before and after digitonin treatment | 1 | Non-parametric one-way ANOVA (Kruskal-Wallis test), P<0.0001 |
| **Figure 9C** FRAP analysis of BiP:sfGFP:HDEL recovery in control and *Atlastin* mutants | Vglut-GAL4;UAS-BiP:sfGFP:HDEL/UAS-RFP:mCD8 (Control)<br><br>Vglut-GAL4;UAS-BiP:sfGFP:HDEL/UAS-RFP:mCD8;atl$^{2}$/atl$^{def}$ (*Atlastin* mutant (Phenotype A))<br><br>Vglut-GAL4;UAS-BiP:sfGFP:HDEL/UAS-RFP:mCD8;atl$^{2}$/atl$^{def}$ (*Atlastin* mutant (Phenotype B)) | 3 larvae, 10 NMJs<br><br>2 larvae, 6 NMJs<br><br>3 larvae, 8 NMJs | Non-linear fit (one-phase association equation) | 1 | Extra sum-of-squares F test |
| **Figure 10B** Endogenous BiP levels in *Atlastin* mutants *SoRa microscope* | Vglut-GAL4 (Control, no marker)<br><br>Vglut-GAL4;;atl$^{2}$/atl$^{def}$ (*Atlastin* mutant, no marker) | 8 larvae, 16 NMJs<br><br>8 larvae, 16 NMJs | (B) Mean intensity of anti-BiP | 1 | Ordinary one-way ANOVA, P<0.0001 |

| Figure/Panel | Genotype (Condition/Label) | N (# larvae, # boutons) | Measurement | # Datasets | Statistical Test(s) |
|---|---|---|---|---|---|
| | Vglut-GAL4;UAS-BiP:sfGFP:HDEL/+ (Control, BiP:sfGFP:HDEL)<br><br>Vglut-GAL4;UAS-BiP:sfGFP:HDEL;atl$^{2}$/atl$^{def}$ (*Atlastin* mutant, BiP:sfGFP:HDEL)<br><br>Vglut-GAL4;;UAS-tdTomato:Sec61β/+ (Control, tdTomato:Sec61β)<br><br>Vglut-GAL4;;UAS-tdTomato:Sec61β/+;atl$^{2}$/atl$^{def}$ (*Atlastin* mutant, tdTomato:Sec61β) | 8 larvae, 17 NMJs<br><br>4 larvae, 9 NMJs<br><br>8 larvae, 15 NMJs<br><br>5 larvae, 9 NMJs | | | |
| **Figure 10G–H** Co-localization of endogenous BiP with tdTomato:Sec61β | Vglut-GAL4;;UAS-tdTomato:Sec61β/+ (Control)<br><br>Vglut-GAL4;;UAS-tdTomato:Sec61β/+;atl$^{2}$/atl$^{def}$ (*Atlastin* mutant) | 3 larvae, 15 NMJs<br><br>4 larvae, 16 NMJs | (G) Mander's correlation coefficient<br>(H) Cytosolic anti-BiP mean intensity | 1 | (G) Unpaired t-test, P<0.05<br>(H) Unpaired t-test, P<0.05 |
| **Figure 11B, D, F, H** MG132 experiments | Vglut-GAL4;UAS-BiP:sfGFP:HDEL/+ (Control)<br><br>Vglut-GAL4;UAS-BiP:sfGFP:HDEL;atl$^{2}$/atl$^{def}$ (*Atlastin* mutant) | Cell bodies:<br>DMSO 6 brains MG132 7 brains<br>Axon bundle:<br>DMSO 7 larvae/14 bundles<br>MG132 8 larvae/16 bundles<br>NMJs:<br>DMSO 7 larvae/14 NMJs<br>MG132 7 larvae/16 NMJs<br><br>Cell bodies:<br>DMSO 8 brains, MG132 7 brains<br>Axon bundle: | (B) α-FK1 mean intensity<br>(D, F, H) BiP:sfGFP:HDEL mean intensity | 1 | (B) Non-parametric one-way ANOVA (Kruskal-Wallis test), P<0.0001<br>(D, F, H) Non-parametric one-way ANOVA (Kruskal-Wallis test), P<0.0001 |

| Figure/Panel | Genotype (Condition/Label) | N (# larvae, # boutons) | Measurement | # Datasets | Statistical Test(s) |
|---|---|---|---|---|---|
| | | DMSO 7 larvae/15 bundles<br>MG132 7 larvae/13 bundles<br>NMJs:<br>DMSO 7 larvae/15 NMJs<br>MG132 7 larvae/13 NMJs | | | |
